# Unlimited genetic switches for cell-type specific manipulation

**DOI:** 10.1101/470443

**Authors:** Jorge Garcia-Marques, Ching-Po Yang, Isabel Espinosa-Medina, Kent Mok, Minoru Koyama, Tzumin Lee

## Abstract

Gaining independent genetic access to discrete cell types is critical to interrogate their biological functions, as well as to deliver precise gene therapy. Transcriptome analyses have allowed us to profile cell populations with extraordinary precision, revealing that cell types are typically defined by a unique combination of genetic markers. Given the lack of adequate tools to target cell types based on multiple markers, most cell types have remained inaccessible to genetic manipulation. Here, we present CaSSA, a platform to create unlimited genetic switches based on CRISPR/Cas9 (Ca) and the DNA repair mechanism known as single-strand annealing (SSA). CaSSA allows engineering of independent genetic switches that each respond to a specific gRNA. Expressing multiple gRNAs in specific patterns enables multiplex cell type-specific manipulations and combinatorial genetic targeting. CaSSA is thus a new genetic tool that conceptually works as an unlimited number of recombinases and will facilitate genetic access to cell types in diverse organisms.

## Introduction

Cell-type specific genetic access is instrumental for cell type characterization, functional analyses, and precise delivery of gene therapy. For instance, it allows us to manipulate genes, use optogenetics or label a particular cell-type so that we can follow its fate and/or isolate it to analyze its transcriptome. In fact, transcriptome analyses have revealed that in order to define most cell types, a combination of multiple gene markers is required (e.g., Zeisel et al., 2015, Rheaume et al., 2018, Li et al., 2017, Han et al., 2018).

Two main approaches have been used to access specific cell types, a reversible and an irreversible approach. Both approaches utilize drivers (specific regulatory sequences to control gene expression) and the expression of responder genes (reporter and/or effector genes). In the reversible approach, the responder is directly controlled by the driver. In this case, the expression of the gene mirrors the driver activity (reviewed in Schenborn and Groskreutz, 1999). However, in the irreversible approach, the driver controls expression of a genetic switch that induces an irreversible change in the DNA, leading to a permanent change in the expression of the responder (Sauer and Henderson, 1988, Golic and Lindquist, 1989, Orban et al., 1992). This approach has traditionally been implemented with site-specific recombinases such as Cre (Lakso et al., 1992) or Flipase (Struhl and Basler, 1993). These enzymes catalyze the DNA recombination between two identical recombinase-specific sites, controlling the expression of a gene by removing a STOP cassette or by reversing the gene’s orientation. Thus, while the reversible approach needs constant activity of the driver for responder gene expression, the irreversible approach requires only transient driver expression in the cell-type of interest.

The irreversible approach has been widely used to target specific cell populations (reviewed in Luan and White, 2007, Del Valle Rodríguez et al., 2012, Luo et al., 2018). This strategy is powerful, in part due to its modular nature, which led to the generation of many transgenic lines with the expression of a recombinase controlled by specific drivers. These lines can be then used to trigger different responder genes in a tissue-specific manner. The collection of responders is enormous, including complex reporters such as Brainbow (Livet et al., 2007), calcium sensors (Zariwala et al., 2012), toxins (Ivanova et al., 2005) or optogenetic effectors (Kätzel et al., 2011). Similarly, these lines can induce conditional mutagenesis when crossed to a line with a specific gene flanked by target sites for the recombinase (reviewed in Lewandoski, 2001). More interestingly, the irreversible approach has also been essential for developmental studies, as one can trigger a reporter in a specific progenitor cell and then trace the subsequent progeny (Buckingham and Meilhac, 2011).

Despite these contributions, the irreversible approach’s ability to access specific cell types is limited by the driver used to express the recombinase. Few genes are truly expressed in only one cell type. Individual cell types are rarely defined by a single marker, but rather by the combination of multiple genes. In recognition of this need, intersectional strategies use two recombinases to express genes only in those cells positive for two different drivers (reviewed in Luan and White, 2007). Yet even this is insufficient considering that 1) multiple genes are necessary to resolve most cell types and 2) the number of recombinases is limited (Nern et al., 2011). This problem is exacerbated when a certain experiment requires targeting two or more cell types at the same time, as occurs in the analysis of neuronal circuits. Moreover, some characterized recombinases are toxic (Loonstra et al., 2001, Forni et al., 2006, Janbandhu et al., 2014) or are not equally effective across different model organisms (Heffner et al., 2012, Hermann et al., 2014). Therefore, even when we know the set of markers defining a cell-type of interest, it may remain genetically inaccessible with current tools.

To address this limitation, we have developed CaSSA, a platform for irreversible genetic access that conceptually works as an unlimited number of recombinases. This system builds upon two universal elements: 1) the specificity of Cas9, an RNA-guided DNA endonuclease associated with the CRISPR system (Gasiunas et al., 2012, Jinek et al., 2012) and 2) single-stranded annealing (SSA), an evolutionarily conserved mechanism for DNA repair (Lin et al., 1984, Ivanov et al., 1996). Here we show that CaSSA can report multiple drivers orthogonally (non-cross reacting) and achieve exceptional specificity by intersecting multiple expression patterns. This study provides proof-of-principle for a methodology that will have general applicability when genetic access to specific cells is required.

## Results

### CaSSA combines Cas9 targeting and SSA-induced DNA editing to switch on a heritable reporter

CaSSA entails a genetic switch responsive to Cas9 induced double strand breaks (DSB). The Cas9 enzyme can introduce a DSB in a DNA region defined by a 20 bp target sequence that is recognized by a guide RNA (gRNA). In practice, this sequence provides the unlimited potential of CaSSA, as there are about 4^^20^ possible unique gRNA targets, and therefore 4^^20^ potential genetic switches that can be triggered. To activate a switch, we took advantage of the DNA repair mechanism known as SSA. This conserved pathway repairs DSBs occurring between two direct repeats by removing one of the repeats and the intervening sequence (reviewed in Bhargava et al, 2016). Since the repair outcome is predictable, we designed a reporter gene that would initially be inactive but would become active after inducing SSA. To do that, the open reading frame of this reporter is interrupted by a switch cassette (Figure 1A). This cassette consists of two direct repeats forming part of the reporter, separated by a specific gRNA target site. After the induction of a DSB by Cas9, the SSA repair reconstitutes a functional reporter. Similar to recombinases such as Cre, this mechanism induces an irreversible change. Therefore, expressing Cas9 and the specific gRNA in neuroblasts triggers the expression of the reporter not only in neuroblasts but also in their progeny.

**Figure 1.**
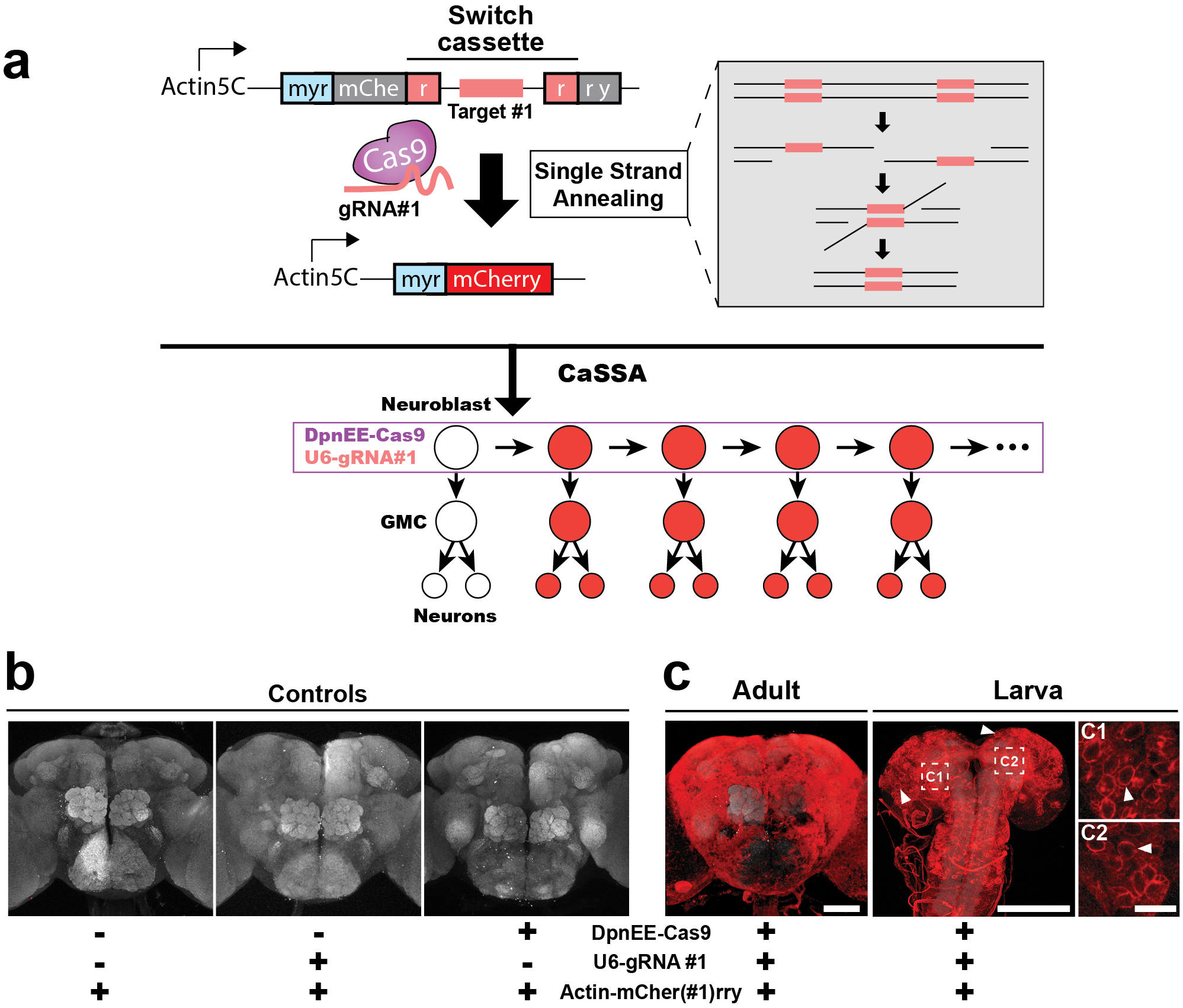
CaSSA combines Cas9 targeting and SSA-induced DNA editing to switch on a heritable reporter. A)Implementation of CaSSA switch reporter activation in Drosophila neuroblasts. CaSSA is based on Cas9 and SSA. An mCherry reporter gene is interrupted with a switch cassette, which consists of a target site specific for the gRNA#1 and two direct repeats (r, part of the mCherry sequence). Cas9 and gRNA#l induce a DSB at the target site of the reporter gene. This is repaired by SSA (grey inset) which reconstitutes and thus activates the reporter gene (red). Cas9 expression is restricted to neuroblasts by the DpnEE promoter (purple outline). Because this change is irreversible, all the progeny derived from that neuroblast will inherit the activated reporter (red). B-C) Maximal confocal projections ofwhole-mount brains areshown (unless otherwisenoted).Red, immunohistochemistry for mCherry. Gray, nc82 counterstain to reveal neuropil. Representative images ofN=l5 brains. Scale bars= 150 µm in C (scale bar in adult panel also applies to B) and 25 µm in C2 (also applies to Cl). B)No red fluorescence is observed in flies missing any of the three transgenes. C)mCherry flourescence is activated in flies bearing all the three CaSSA components: reporter, Cas9 and gRNA. In the larval brain, most neuroblasts (arrowheads) express the mCherry functional protein. The activated switch is inherited by the neuroblast progeny leading to ubiquitous red fluorescence in the adult. Cl) and C2) High magnifications from a single focal plane in C) showing characteristic neuroblast morphology.

To evaluate the efficiency of CaSSA in Drosophila, we generated three transgenes: (1) Actin5C-mCher(#1)rry, an Actin5C promoter driving a conditional mCherry reporter containing a switch cassette consisting of two direct repeats (400 bp of the reporter sequence) separated by a target site specific for the gRNA#1, (2) DpnEE-Cas9, Cas9 under a neuroblast-specific deadpan (Dpn) driver that is active in all neuroblasts outside of the optic lobe and is silent in ganglion-mother cells (GMC) and post-mitotic neurons (Awasaki et al., 2014) and (3) U6-gRNA#1, gRNA #1 driven by the ubiquitous U6 promoter. Neither the U6-gRNA#1 nor the DpnEE-Cas9 transgenes alone were able to trigger the reporter (Figure 1B). Only after combining all three elements, we observed the expression of the reporter (Figure 1C). In larvae, this reporter was expressed in most neuroblasts and their progeny (Figure 1C1&C2). This led to an apparent ubiquitous expression of the reporter in the adult brain.

To confirm the efficacy of CaSSA in Drosophila, we set out to test individual repair events (Figure S1). To do that, we set up crosses to combine the three CaSSA elements (Actin5C-Cas9, U6-gRNA#1 and Actin5C-mCher(#1)rry) in the same flies (G0). We then established independent crosses between the G0 and double balancer flies and selected a single fly bearing the reporter from each cross. While the G0 fly is a mosaic, all the cells in the G1 contain the same allele for the repaired reporter. This procedure guaranteed that each fly reflected a single repair event, showing that 96 out of 100 repair events yielded a functional reporter. To analyze the repair outcome in the 4 cases where this repair seemed to fail, we attempted to sequence the reporter. In all cases we failed to amplify the reporter, suggesting that large deletions occurred that prevented PCR amplification (Kosicki et al., 2018).

### Multiple gene patterns can be orthogonally reported by CaSSA

After testing the feasibility of CaSSA to trigger a reporter gene, we set out to prove the ability of this system to report lineage-specific drivers expressed in neuroblasts. This should immortalize the pattern of these drivers into the progeny generated from that neuroblast (Figure 2A). Cell-type specific genes are transcribed from type II promoters. However, these promoters are not appropriate for gRNA production, as the mRNA produced contains additional sequences that may hinder the gRNA function (Gao and Zhao, 2014). To use type II promoters to generate gRNAs with no extra sequence, we had to flank these gRNAs with the hammerhead (HH) and hepatitis delta virus (HDV) ribozymes. Given their self-cleaving activity, these ribozymes are sufficient to process a functional gRNA from a mRNA in the absence of any other cofactor (Gao and Zhao, 2014). To increase the efficiency of this system, we initially used three copies of these ribozyme/gRNA cassettes to generate a transgenic line with the antennal lobe (AL) driver 44F03 (Jenett et al., 2012, Awasaki et al., 2014) driving the expression of the gRNA#1. This driver is expressed in all four AL lineages (ALad1, ALl1, ALv1, and ALv2), with some minor activity in the mushroom body (MB) and ellipsoid body (EB) lineages. We generated another line with the gRNA#2 expressed under the regulation of the 41A10 driver (Jenett et al., 2012, Yang et al., 2016, Figure 2B). This is active in MB lineages, with some minor activity in other seemingly random lineages in the brain. After crossing these pol II driven, ribozyme flanked gRNA lines to a line with DpnEE-Cas9, Actin5C-mCher(#1)rry and Actin5C-EGF(#2)FP, we found patterns very similar to those obtained with a classical recombinase (Awasaki et al., 2014, Yang et al., 2016). Each gRNA was specific for its reporter, which allowed us to report both drivers without any cross-talk.

**Figure 2.**
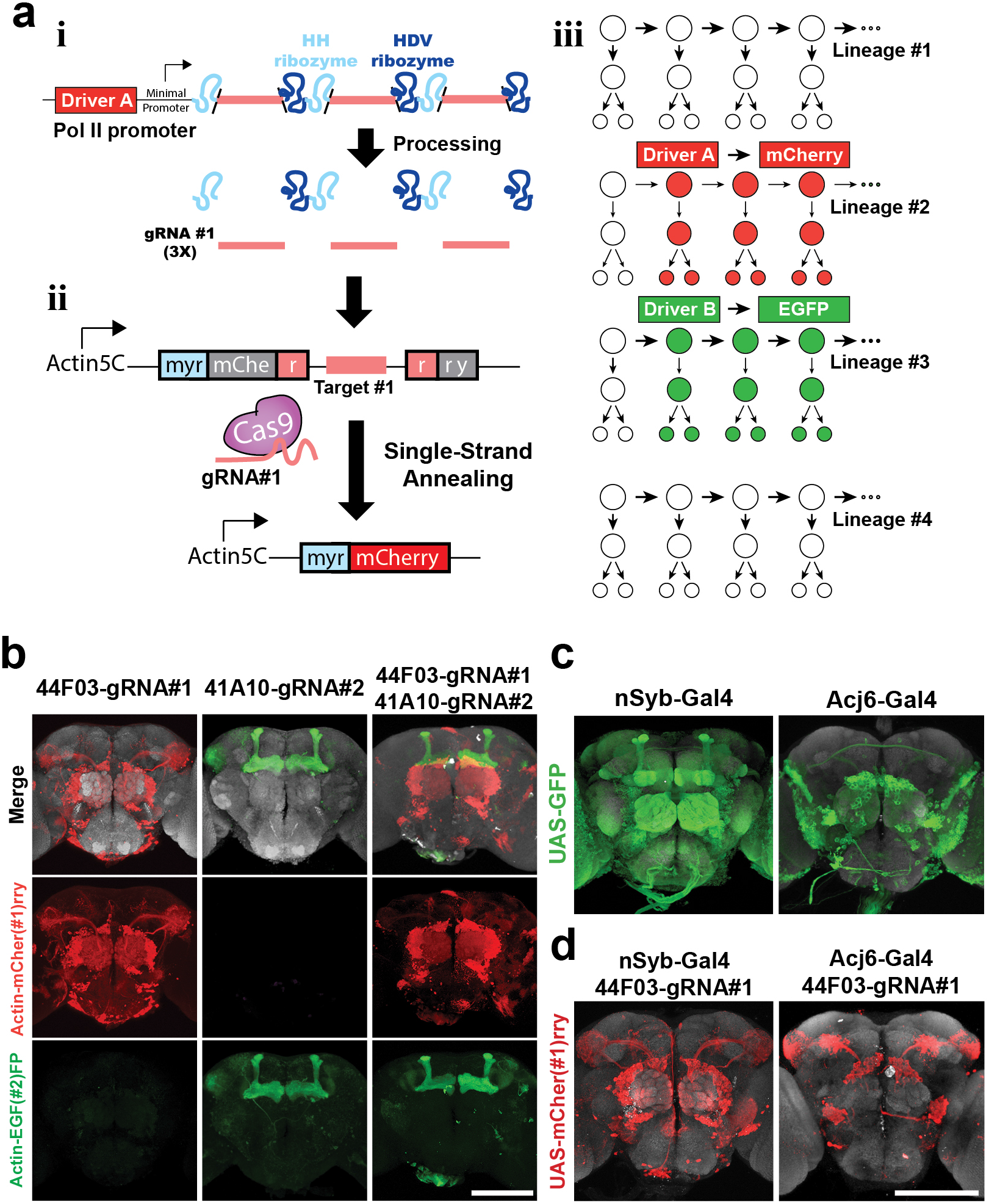
Multiple gene patterns can be orthogonally reported by CaSSA. A) Scheme illustrating the simultaneous use of multiple CaSSA switch reporters in the brain. i) Tissue specific induction of gRNA is accomplished by utilizing pol II promoter, consisting of tissue specific enhancer/driver element (Driver and a minimal promoter. gRNAs are not properly processed when transcribed by RNA polymerase II, therefore we flanked three copies of gRNA (pink) with ribozymes (blue). Ribozymes process the mRNA producing gRNA with no extra sequence. ii) gRNAs together with Cas9 induce a DSB at the target site. Repair via SSA reconstitutes the reporter gene. iii) Different tissue specific drivers can be paired with switch reporters and neuroblast specific Cas9 to label lineages of interest. B)Drivers 44F03 and 41Al0 were reported simultaneously without any crosstalk by using two different gRNAs, each one specific for a different reporter. C)Expression patterns of nSyb andAcj6 using the Gal4/UAS system. D)Intersection between CaSSA and the Gal4/UAS system. Examples for nSyb-Gal4 andAcj6-Gal4 patterns intersected by 44F03-gRNA#l. Maximal projection pictures of whole-mount brains are shown. Red, immunohistochemistry for mCherry. Green, immunohistochemistry for EGFP. Gray, nc82 counterstain. Representative images of N=20 brains. Scale bars= 150 µm in B and D (also applies to C).

To extend the application of this system, we also created UAS driven reporters. In this way, CaSSA could be used to intersect with the Gal4 system. To test this idea, we used 44F03-gRNA#1 to refine the pan-neuronal pattern of nSyb-Gal4 (Figure 2C). As nSyb-Gal4 is expressed in most of the mature neurons in the brain (Pfeiffer et al., 2008), this intersection showed the 44F03-gRNA#1 pattern (Figure 2D). We also used 44F03-gRNA#1 to intersect the Acj6-Gal4 driver, which is expressed in olfactory sensory neurons, projection neurons generated from the adPN and ALl1 lineages and some other neurons in the lateral part of the brain (Bourbon et al., 2002, Lai et al., 2008; Figure 2C). This intersection restricted the Acj6-Gal4 pattern to only those neurons generated from AL lineages (Figure 2D).

### CaSSA can be used as a powerful gene trap system to capture endogenous transcription

Gene traps provide a formidable tool to identify interesting gene expression patterns (O’Kane and Gehring, 1987). This relies on inserting a reporter gene randomly into the genome, thus capturing the expression of the gene at the site of insertion. When this expression pattern is restricted to a cell type of interest, such a tool may be valuable for further applications. Yet, the current method requires the insertion to occur in the right orientation and in the same open reading frame as the endogenous gene, which greatly reduces the probability of capturing an endogenous expression pattern. We reasoned that CaSSA would constitute a very powerful gene trap system, as the gRNA only requires transcription to report the gene expression pattern. To show proof-of-principle on the application of CaSSA to generate useful drivers, we devised a new gene trap construct. By modifying the design used in the MiMIC system (Venken et al., 2011), we generated a construct with two gRNAs (three copies flanked by ribozymes) in opposite orientations so that we could report both sense and antisense transcription (Figure 3A). To identify each insertion, we also included a *yellow*^+^ dominant body-color marker, located between the gRNAs. This cassette was flanked by two inverted ΦC31 attP sites, which allows us to replace this construct with any other DNA (Bateman et al., 2006). We introduced the full construct between inverted repeats for the Minos and P-element transposons so that we could use either of the corresponding transposases to mobilize the reporter.

**Figure 3.**
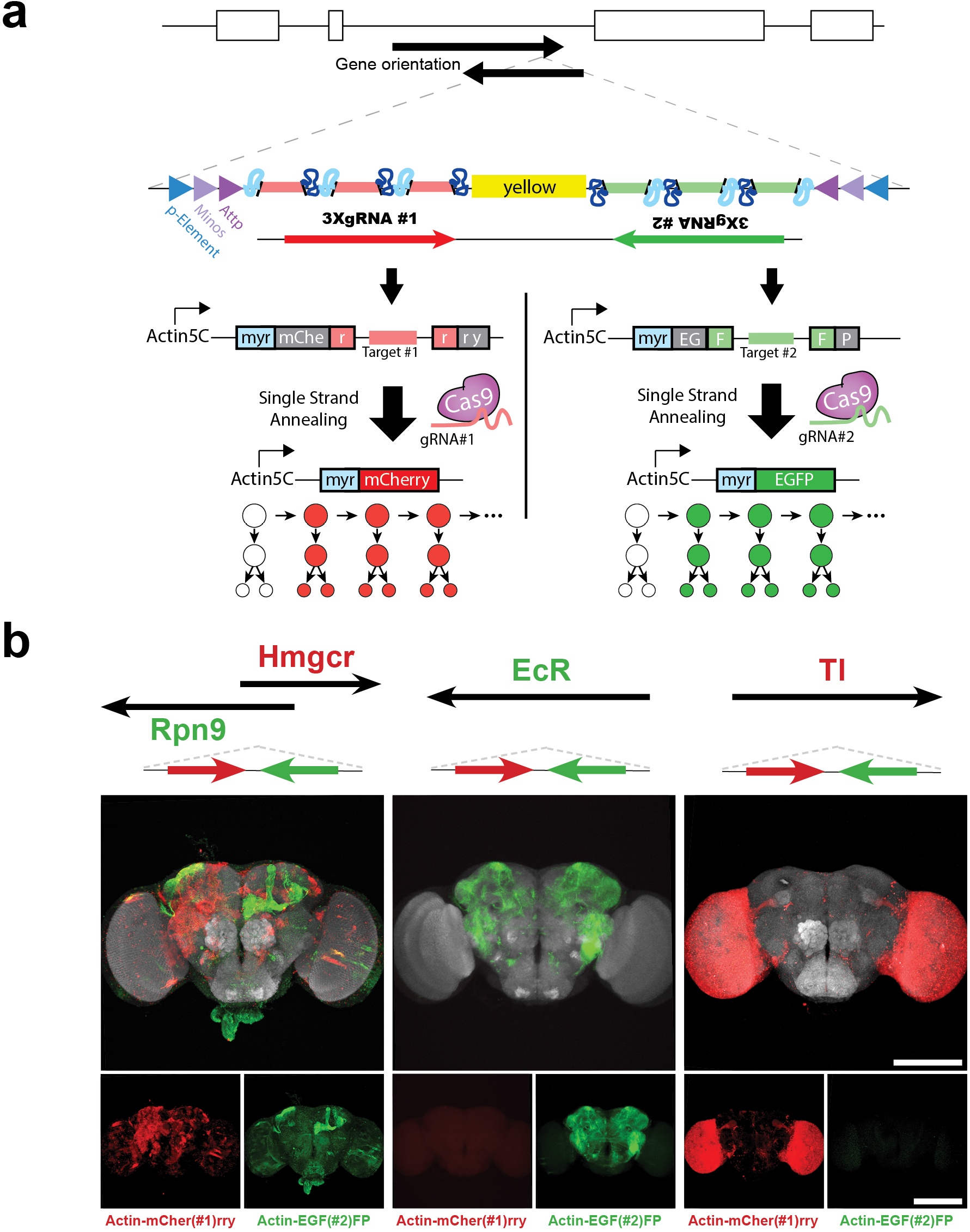
CaSSA can be used as a powerful gene trap system to capture endogenous transcription. A)Scheme of the gene trap construct implemented with CaSSA. Three copies of two different gRNAs (red gRNA#1, green gRNA#2)/ribozyme (blue) cassettes were cloned in opposite directions. To select positive lines, a yellow+ marker was inserted between cassettes. With this design, each gRNA orthogonally reports a different strand (sense or antisense) by activating a different reporter. B)Examples of three lines with a labeling pattern that reflects the orientation and position of the construct into the genome. These lines were crossed to flies expressing Actin5C-Cas9, Actin5C-mCher(# l)rry andActin5C-EGF(#2)FP. Maximal projection pictures of whole-mount brains are shown. Red, immunohistochemistry for mCherry. Green, immunohistochemistry for EGFP. Gray, nc82 counterstain. Scale bars= 150 µm. See also Figure S2.

To demonstrate the versatility of CaSSA as a gene trap, we first inserted this construct by using the P-element transposase, generating 55 different yellow + lines. Then, we selected 7 starting lines with the transposon located on the X chromosome to mobilize with the Minos transposase, generating 115 additional lines (Figure S2, Table S2). Subsequently, we crossed each of these lines to a line expressing Actin5C-Cas9, Actin5C-mCher(#1)rry and Actin5C-EGF(#2)FP. For pre-screening, we chose Actin5C-Cas9 to find any interesting patterns regardless of the cell origin or identity. Any interesting insertion with a restricted and reliable pattern would then be crossed to DpnEE-Cas9 to find useful drivers in neuronal lineages. Out of the 170 lines, 146 (85.9%) showed some reporter expression, confirming the efficiency of CaSSA as a gene trap system. Each line produced a distinct expression pattern ranging from widespread (scored as 5, Table S2) to restricted (scored as 1, Table S2). There was a bias towards lines reporting only the gRNA#2 (64/170; 37.6%) compared to those reporting only the gRNA#1 (27/170; 15.9%). This could have arisen from the asymmetry introduced in the construct by the *yellow*^+^ marker, which could favor this orientation in the transposition event. The remaining lines reported both gRNAs (55; 32.4%), confirming the construct could simultaneously capture both sense and antisense transcription. These double trap lines included insertions with widespread (i.e. F25M03, Table S2) as well as restricted patterns (i.e. J001, Table S2).

To show the relation between transcription and the reporter pattern, we performed inverse PCR to map the insertion site for 14 lines. In nine out of 14 lines, the observed labeling reflects the transcription described in the genomic map (Examples in Figure 3B, Table S2; Flybase). Three out of 14 lines showed expression of both reporters despite being integrated into a single annotated gene, which might reflect some residual transcription not previously reported. The remaining two lines showed no reporter expression despite the insertion’s location in an annotated gene. These lines correspond to insertions in CadN2 and CG30438, both genes having low expression in the central nervous system (RNA-seq-CNS and adult head track; Flybase). In conclusion, CaSSA was able to report transcription in 86% of the lines generated, revealing its potential as a gene trap system. However, most of our lines showed either a stable but ubiquitous or a rather restricted but variable pattern.

### Increasing the number of gRNA copies improves the targeting effectiveness

As we did not find lineage specific genes, we reasoned that our gene trap system might be unable to report potentially interesting genes due to a lower transcription level. If true, increasing the number of gRNA copies should help capture transcription from these genes. In an effort to address the influence of copy number in the expression pattern, we went back to the well-characterized 44F03 pattern. We generated three additional transgenic lines expressing 6, 12 or 24 gRNA copies. As shown in Figure S3, increasing the number of copies also increased the percentage of hemibrains labeling a particular lineage. This was more evident for lineages such as EB or MB, which suggests that the 44F03 transcriptional activity is weaker in these lineages. Interestingly, the probability of hitting some lineages such as the lateral AL lineage was higher than 90%, confirming that CaSSA is very effective, not only in the germ line, but also in neuroblasts. We also found that for most lineages, the targeting effectiveness was saturated with 12 copies of gRNA.

After addressing the effect of gRNA copy number on the labeling effectiveness, we set out to test this idea for an endogenous homeobox gene. We selected Dr as it is involved in dorsoventral patterning and neuroblast specification (D’Alessio and Frasch, 1996, Isshiki et al., 1997). To target Dr we induced recombinase-mediated cassette exchange (RMCE) in an existing MiMIC line (Venken et al., 2011; Figure 5A). In parallel, we generated two transgenic lines with cassettes expressing 3 or 12 copies of the gRNA#1. Each of these lines also expressed the gRNA#2 in reverse orientation to report any potential antisense transcription. Subsequently, we crossed each of these lines to flies expressing Actin5C-Cas9, Actin5C-mCher(#1)rry and Actin5C-EGF(#2)FP. We chose Actin5C-Cas9 rather than DpnEE-Cas9 so that we could capture any early expression, as it is known that Dr expression is first observed before neuroblast delamination (D’Alessio and Frasch, 1996). No significant expression was found for the gRNA#2, confirming the absence of antisense transcription in the genome map (Flybase). For the gRNA#1, both lines showed most of the reporter expression in lateral horn (LH) lineages, and less frequently in the AL (Figure 5B, C). The 3-copies line reported expression in the LHl1, SLPal1, VLPl4, VLPp1, ALlv1 and ALv2 lineages (Yu et al., 2013). Additionally, the 12-copies line also showed expression for ALad1 and ALl1. These lineages corresponded to those labeled by using our previous methodology based on driver immortalization with recombinases (Awasaki et al., 2014), although this methodology also reported the VPNp1, VESa1 and PSa1 lineages (Figure 4S). Despite 3 copies was enough to label most of the lineages reported with 12 copies, increasing the copy number resulted in a drastic improvement of the targeting effectiveness for all lineages. Interestingly, both lines with 3 or 12 copies were perfectly viable as a homozygous, in contrast to the line with the KD recombinase inserted in the same locus. This suggests that expressing multiple gRNAs has no deleterious effects and does not interrupt the expression of the endogenous gene.

**Figure 4.**
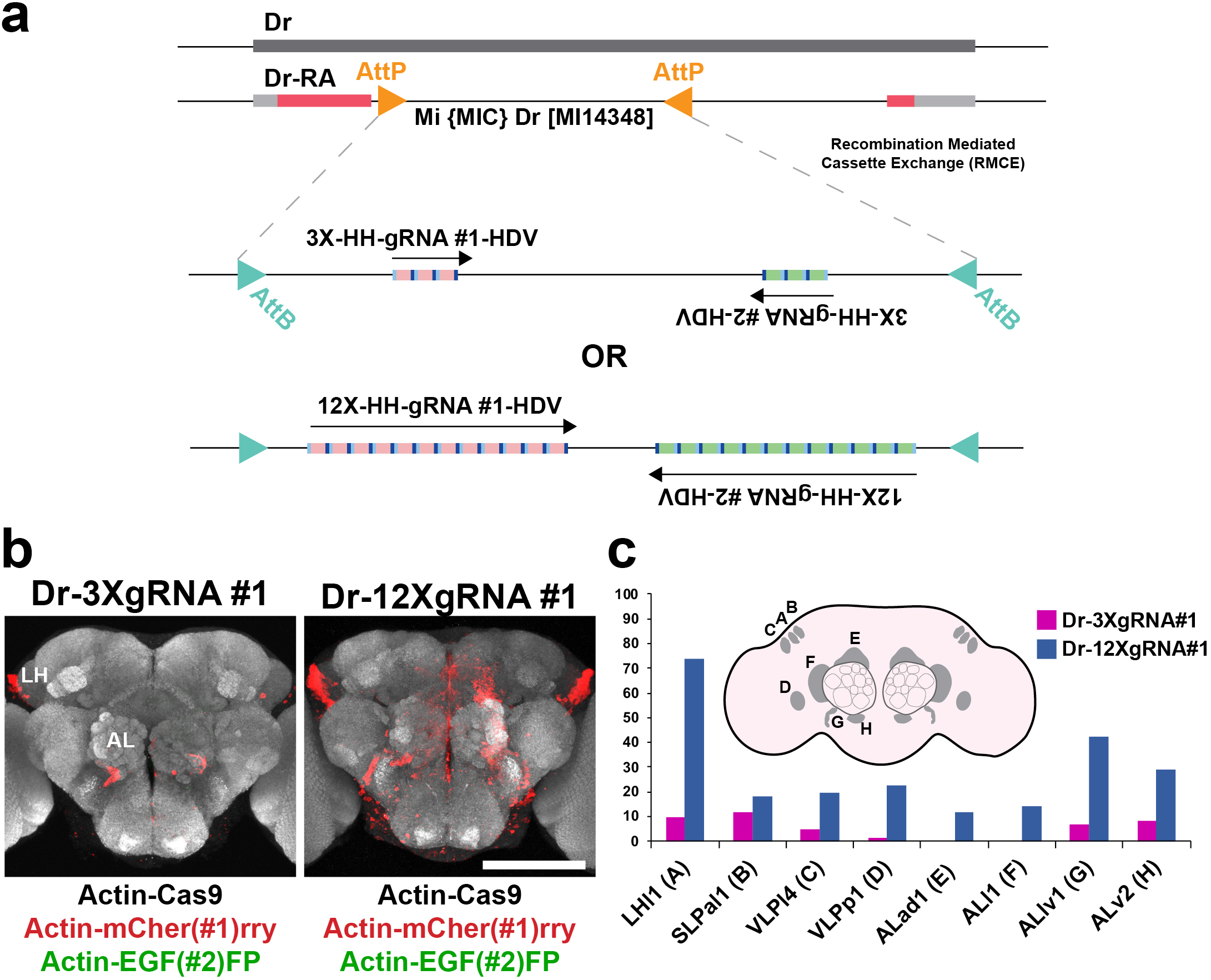
Increasing the number of gRNA copies improves the targeting effectiveness. A)Illustration showing the strategy to target Dr based on a MiMIC line. Two different transgenic lines were generated by exchanging the AttP-flanked genomic region in the MiMIC line for a cassette previously flanked by AttB sites (RMCE). Each cassette contained either 3 or 12 copies of ribozyme flanked gRNAs. Two different gRNAs were used to report sense and antisense transcription. B)Three copies of gRNA are sufficient to report the Dr expression pattern. Flies with 3 or 12 copies of gRNA inserted in the Dr locus were crossed to a line bearing Actin5C-Cas9, Actin5C-mCher(#l)rry and Actin5C-EGF(#2)FP. No significant expression was found for the antisense gRNA (green reporter). A similar qualitative pattern was found for 3 or 12 gRNA copies. C)12 copies of gRNA resulted in a drastic increase of targeting effectiveness. The position of the soma for each lineage is shown in the cartoon. Data are represented as the percentage of hemibrains (N=60) with reporter expression for each lineage. Maximal projection pictures of whole-mount brains are shown. Red, immunohistochemistry for mCherry. Green, immunohistochemistry for EGFP. Gray, nc82 counterstain. Representative images of N=30 brains. HH=Hammerhead ribozyme; HDV=HDV ribozyme. Scale bar= 150 µm in B. See also Figures S3, S4.

**Figure 5.**
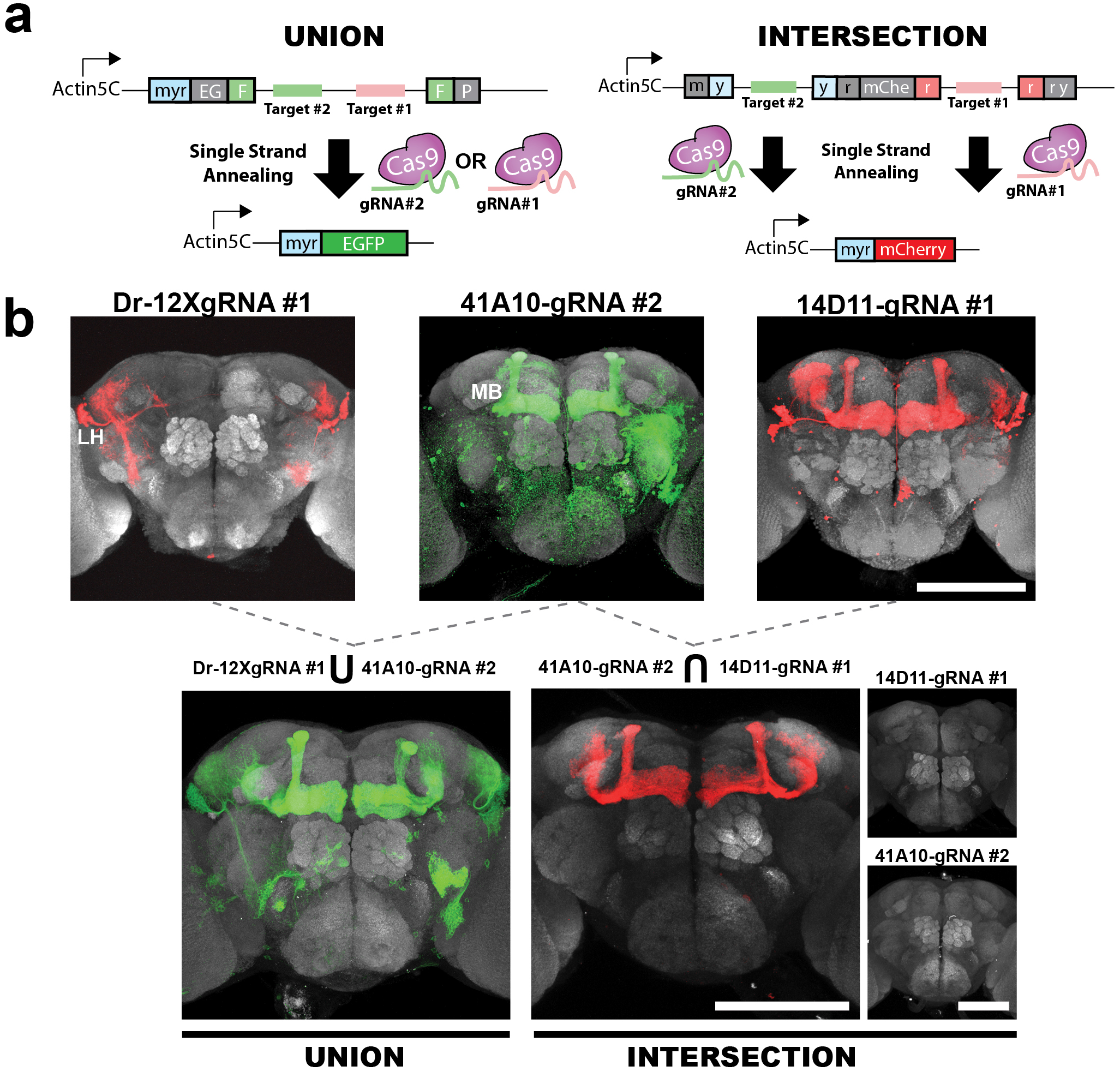
Generating additional tools for enhanced accessibility: union and intersection. A)Schematic design for the Union and Intersection reporters. For the Union construct, two target sites for different gRNAs were placed between the direct repeats (green box, F). Providing either gRNA would trigger SSA and reconstitute the reporter. For the Intersection construct, two different switch cassettes were placed along the reporter’s open reading frame. Since each cassette is triggered by different gRNAs, both of them are required to reconstitute a functional reporter. B)Proof-of-principle for the Union and Intersection constructs. Top panels show individual driver/reporter patterns for comparison. For the Union experiment, providing two gRNAs results in the combination of Dr (lateral horn lineages) and 41Al0 (MB lineages) patterns. In the Intersection, only lineages that express both 41Al0 and 14Dll are labeled (MB lineages). No fluorescence exists when only a single gRNA is expressed, both gRNAs are required to activate the intersection reporter. In all cases DpnEE-Cas9 was used. Maximal projection pictures of whole-mount brains are shown. Red, immunohistochemistry for mCherry. Green, immunohistochemistry for EGFP. Gray, nc82 counterstain. Scale bars= 150 µm.

### Generating additional tools for enhanced accessibility: union and intersection

Given the specificity of Cas9, multiple gRNAs can be simultaneously utilized. We therefore exploited concurrent employment of multiple gRNA variants to target neuronal lineages. We reasoned that implementing the boolean gates OR and AND could be very useful. The OR gate could make targeting a particular lineage more reliable if one has multiple weak or noisy drivers. Also, an OR gate could be used to target a full population of cells by using several drivers specific to different subpopulations. Similarly, the AND gate could help gain specificity by labeling lineages that express a combination of two or more genetic markers.

To implement the OR gate (Union) we designed a single switch with target sites for two different gRNAs, gRNA #1 and #2 (Figure 5A). To demonstrate the utility of this reporter, we expressed gRNA #1 driven by the Dr endogenous regulatory sequences and gRNA #2 driven by the MB driver 41A10 (Figure 5B). As shown above, Dr-gRNA#1 consistently labels lineages in the LH and occasionally the AL but never the MB. In contrast, 41A10 mostly labels MB lineages together with some other low-frequency lineages, but very rarely the LH. After inducing the union reporter, we obtained a combined pattern comprising the MB and some LH lineages (Figure 5B).

To generate the AND gate (Intersection) we designed another reporter construct with two switch cassettes: one in the myristoylation signal sequence (with the target site #2) and the other in the mCherry sequence (with a target site for the gRNA#1) (Figure 5A). To test this construct, we used 41A10 for the gRNA#2 and 14D11 for the gRNA#1. Both drivers label MB (composed by 4 identical clonal units), as well as some other non-coincident lineages. After introducing both constructs together with Cas9, we found labeling exclusively in the MB (Figure 5B). In contrast, expressing only one gRNA was not sufficient to induce any reporter expression. As the intersection involves two DSBs, we predicted that the occurrence of inter-target deletions could impede the expression of the reporter gene. Another factor that might affect the efficiency of this construct is the probability for each driver to label the same MB clonal unit, as they do not label all the clonal units in every brain. Therefore, to analyze the efficiency of this construct, we first estimated the probability of double labeling the same clonal unit when each driver triggered an independent reporter. By using EGFP to report 41A10 and mCherry as a reporter of 14D11, we found double reporter labeling in 14.8% of clonal units (14 out of 96 clonal units, 24 brains). This allowed us to estimate the efficiency of the intersection construct at ~40% as it reported 39.6% of these double events (6 out of 104 clonal units, 26 brains).

### CaSSA is effective in a model vertebrate

To increase the general applicability of CaSSA, we wanted to prove its effectiveness in vertebrates. We reasoned this should be possible since both Cas9 (reviewed in Doudna and Charpentier, 2014) and SSA (reviewed in Bhargava et al, 2016) work efficiently across different organisms. To test CaSSA’s feasibility in zebrafish as a model vertebrate, we designed a plasmid encoding the following elements in a single open reading frame: 1) a positive control for transfection (mCherry), a CaSSA reporter for the gRNA#1 and 3) Cas9. The three elements were driven by the ubiquitous promoter Ubi (Figure 6A). In the same construct, we either introduced a non-matching gRNA (#3) or the corresponding gRNA#1 under the U6 promoter. This plasmid also contained the Tol2 transposon repeats, for the genomic integration of this construct in the transfected cells (Don et al., 2017). After injecting zebrafish embryos with mRNA for the Tol2 transposase and the CaSSA plasmid with non-matching gRNA (negative control) only a few mCherry positive cells showed expression of the reconstituted reporter (1.3% +/-0.9; Figure 6B). In contrast, injecting Tol2 mRNA and the plasmid with the correct gRNA #1 produced a 100-fold increase in reporter expression, with virtually all the mCherry positive cells expressing the second green reporter (99% +/-0.5).

**Figure 6.**
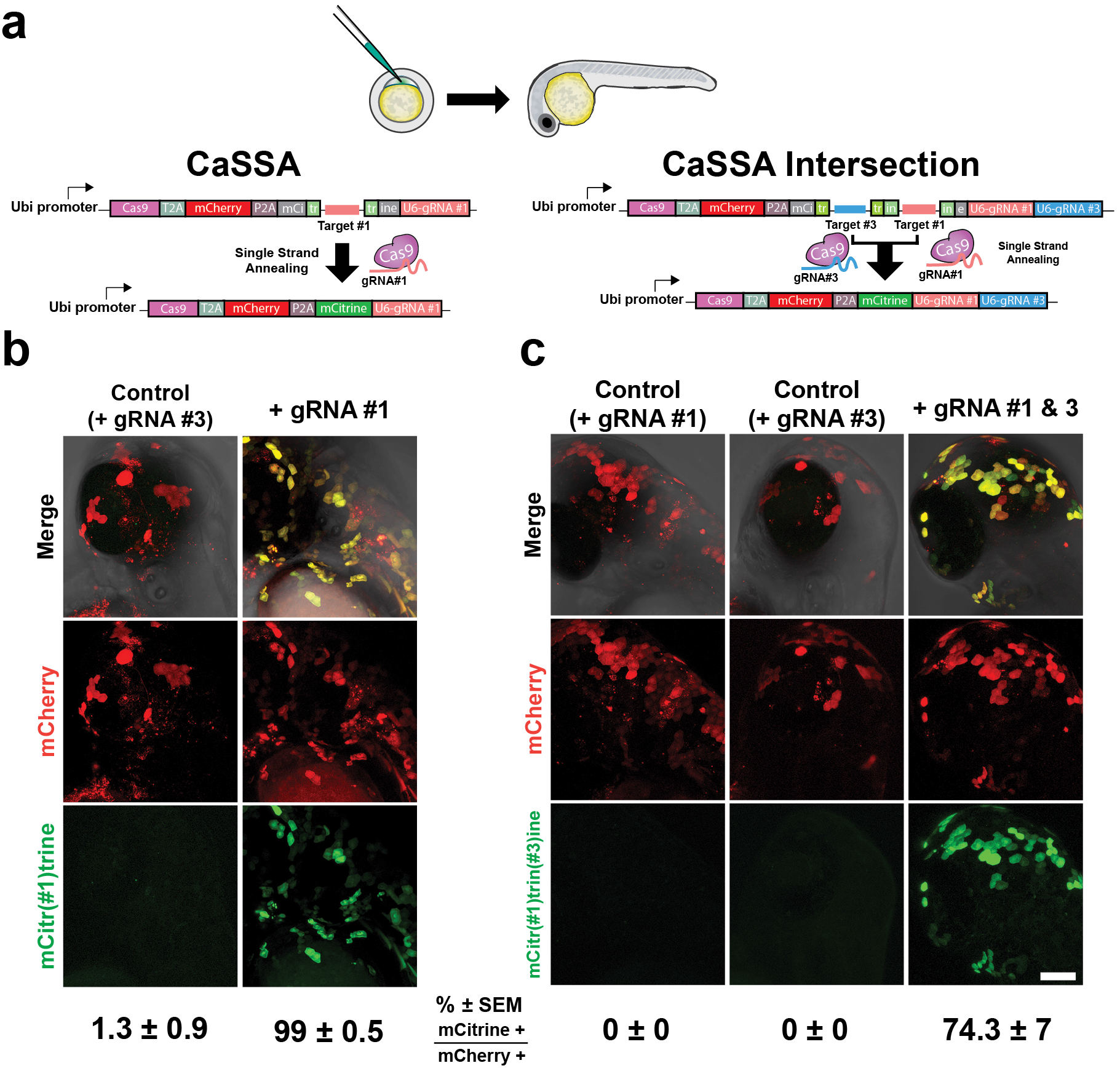
CaSSA is effective in a model vertebrate. A)Cartoon illustrating the experimental procedure and the constructs design. Each construct contains Tol2 inverted repeats for genomic integration. Since not all cells capture/integrate these constructs, a transfection marker (mCherry) is encoded in the same open reading frame (separated by T2A or P2A peptides) as Cas9 and the switchable reporter. This cassette is under regulation of the ubiquitous promoter Ubi. Downstream of this cassette we placed the corresponding gRNA(s) driven by an U6 promoter. B)CaSSA performance in fish. In the control, the construct has a mismatching gRNA and results in almost no cells expressing mCitrine. However, in the experimental construct that contains the matching gRNA #1, virtually every mCherry positive cell expressed the reconstituted mCitrine reporter. C)CaSSA intersection in fish. In the control scenarios, providing only one of the gRNAs with targets in the intersection construct did not trigger reporter fluorescence in any cell. Only after providing both gRNAs was it possible to see the reconstituted reporter in about 3 out of 4 cells. Maximal projection pictures of whole-mount fish (head region, 2 dpf) are shown. Red, endogenous mCherry signal. Green, endogenous mCitrine signal. Merge is shown with bright field to reveal larval anatomy. Data are represented as percentage (± SEM) of mCitrine positive out of the total number of transfected cells (mCherry positive). Representative images and quantification from at least 5 fish. Scale bar= 100 µm in C (also applies to B).

We showed in flies that the intersection of different gRNAs could label a cell type with an extraordinary level of specificity. To test this idea in zebrafish we generated a CaSSA reporter with two switch cassettes (Figure 6A), similar to the intersection construct shown above (Figure 5A). After injecting only one of the gRNAs, either the gRNA#1 or the gRNA#3 driven by the U6 promoter, we did not find any cell expressing the green reporter (Figure 6C). However, the injection of both gRNAs resulted in the expression of the green reporter in most of the mCherry positive cells (74.3% +/-7). These results demonstrate the effectiveness of CaSSA in a model vertebrate, suggesting this tool could also be applied to diverse models.

## Discussion

CaSSA is a new platform for gaining genetic access into specific cell types. We developed and optimized this system, proving its efficacy in an invertebrate and a vertebrate model. This tool allows one to generate and report multiple drivers, which can be then combined to create complex patterns or intersected to target specific cell types.

Irreversible methods for gaining access into specific cell types have proven extremely helpful, particularly in developmental studies. Taking into consideration that transcriptional diversity may peak during development (Li et al., 2017), targeting differentiated cell types could also be more achievable by using irreversible genetic switches than with dynamic drivers. Traditional recombinases have been widely used as the reference method for irreversible genetic access (Lakso et al., 1992, Struhl and Basler, 1993). However, if we want to take advantage of all the information that transcriptomic data provides to gain access into specific cell types, the number of conventional recombinases fall extremely short. For instance, projection neurons in the AL can only be distinguished using a combinatorial code of 11 genes (Li et. al, 2017). Even if we could find new recombinases, expressing several in the same cell could have deleterious effects (Loonstra et al., 2001, Forni et al., 2006, Janbandhu et al., 2014). CaSSA aims to fill that gap by harnessing the unlimited specificity of RNA-guided CRISPR endonucleases. The other element of this system is SSA, a conserved mechanism for DNA repair in regions with direct repeats (Lin et al., 1984, Ivanov et al., 1996). In contrast to other DNA repair mechanisms such as non-homologous end joining or homology directed repair, SSA has remained overlooked for most biotechnology applications. While reporters utilizing SSA were developed before (Pierce et al., 1999), the lack of Cas9 restrained their scope to studying DNA repair. CaSSA leverages Cas9 to exploit this repair mechanism and create an alternative to traditional recombinases. Besides performing similarly to recombinases, CaSSA offers four big advantages: (1) a virtually unlimited specificity, (2) high efficacy across different models, (3) scarless gene reconstitution and (4) minimal toxicity, as evidenced by the fact that Actin5C-Cas9 (ubiquitous) is homozygous viable, as well as flies expressing up to 6 different gRNAs and DpnEE-Cas9 (data not shown).

An unlimited number of genetic switches does not provide much of an advantage if we lack specific drivers to target our cells of interest. Here we show the versatility of CaSSA applied to search for drivers in an efficient way. As opposed to other gene trap systems based on reporter genes that need to be translated (Venken et al., 2011), CaSSA can report any transcription, regardless of the orientation, open reading frame or insertion site in the gene. Such minimal requirements make it a very powerful gene trap system for seeking new gene patterns in an unbiased manner. Most of the gene trap systems are based on a reporter gene being controlled by the regulatory regions where it is inserted (reversible approach). In contrast, CaSSA or any gene trap based on recombinases (Bohm et al., 2010) do not aim to describe the real-time expression pattern of the gene where they are inserted. Instead, these gene traps elucidate any instance of gene expression occurring during any time point.

Given our interest in Drosophila neurogenesis, we oriented our initial gene trap effort to find patterns restricted to subsets of neuronal lineages. Unfortunately, we failed to find any stable pattern. This is likely due to (1) an insufficient number of lines generated or (2) spatial segregation genes may be low expressed, and thus missed by the gene trap. To test the latter possibility and assuming that similar genes might work at similar transcription levels, we wanted to prove that we could detect this type of genes. We selected Dr as an example, as it is a homeodomain transcription factor involved in neuroblast specification (D’Alessio and Frasch, 1996, Isshiki et al., 1997). We thus confirmed that three gRNA copies were sufficient to report part of the lineages targeted by recombinases. Even if the efficiency is low, this is potentially a valuable reagent as we usually lack drivers to target only few lineages. In fact, increasing the copy number seems to be a good balance between efficiency and specificity. By expressing 12 copies of gRNA the targeting effectiveness increased up to 13-fold (for VLPp1, Figure S4) and only two additional lineages were reported. If necessary, the concentration of gRNA could be further increased in the future by using alternative ways to process the gRNA from the mRNA, such as a tRNA system (Port and Bullock, 2016). Interestingly, despite the embryonic lethality of Dr mutants (Isshiki et al., 1997), our lines containing 12 ribozyme flanked gRNA copies inserted into a Dr intron were homozygous viable. This suggests that splicing precedes ribozyme processing so that the mRNA remains intact. This would make it possible to report heterozygous lethal genes. In contrast, the expression of recombinases requires polyA signals that interrupt the transcription of the endogenous gene.

One of the greatest advantages of CaSSA is the possibility of implementing Boolean gates with multiple gene patterns. In particular, CaSSA permits multi-gene intersections, which can be applied to target specific cell types based on a combination of gene markers. In fact, CaSSA is already a double intersection between two drivers, one for the expression of Cas9 and another one for the gRNA. Also, it allows further intersections given its compatibility with traditional reagents such as the Gal4 system. Moreover, we show proof-of-principle for triple intersection to label cells expressing three different drivers. Despite reduced efficiency, this reporter could be very useful to applications where specificity is a priority over efficiency, such as isolating specific cell types for genome-wide molecular profiling. Notably, a similar intersectional reporter responded very efficiently in zebrafish. Based on the nature of transient Tol2 transgenesis, this could be a result of integration of multiple copies of the transgene into the genome. This therefore suggests a way to improve the efficiency in fly, by inserting multiple copies of the reporter in different landing sites. Similarly, the efficiency of this reporter could be improved by reducing the inter-target deletions. Given the fact that larger deletions are less likely to happen (Liu et al., 2016, Canver et al., 2014), we could design a new reporter with long introns where the switch cassettes would be far from each other. Also, a single DSB could be enough to achieve triple intersections by using the wild type CRISPR system where the gRNA is split into two parts: crRNA and tracrRNA. In this way we could activate a reporter by inducing a single DSB only in those cells expressing each of the three elements. Yet another possibility would be using a variant of Cas9 to activate a reporter by single-base editing (Komor et al., 2016), rather than inducing DSBs in the DNA.

Besides its application to intersect multiple gene patterns, the fact that CaSSA works as an unlimited (scarless) recombinase holds immense potential. It allows us to target multiple cell populations simultaneously, which would be very useful to study neuronal circuits. Also, we envisage that this system could work as an integrative solution to consolidate other genetic tools. For instance, we can generate a reporter with multiple sites for a specific gRNA, upstream from a minimal promoter. Thus, we could activate the reporter by using a dead Cas9 variant (with no nuclease activity) fused to a transcriptional activator domain (Ewen-Campen et al., 2017). We can multiplex them with diverse reporter genes responding to distinct gRNAs. In this way, we could barcode multiple genes by integrating different gRNAs and using the gene-specific gRNAs to report the real-time expression of the targeted genes (reversible approach) or activate a reporter in an irreversible way to label a whole lineage. In conclusion, CaSSA lays the groundwork for the genetic access of the future by unleashing the unlimited specificity of RNA guided CRISPR endonucleases.

## Acknowledgments

We thank Haluk Lacin and all members of Tzumin’s lab for their comments and feedback, especially Rosa Miyares for critical reading and input on the manuscript. We thank Denise Montell for her feedback on the manuscript. We thank Janelia Fly Core and Molecular Biology shared resources, especially Xiaorong Zhang for her excellent technical support. We thank our suppliers Rainbow, Genscript and Benchling for their services. We thank C. Sullivan for administrative support. This work was supported by Howard Hughes Medical Institute.

## Author contributions

Conceptualization, J.G.-M. and T.L.; Methodology, J.G.-M. and T.L.; Investigation, J.G.-M., C.-P.Y., I.E.-M. and K.M.; Writing – Original Draft, J.G.-M. and T.L.; Writing - Review & Editing, J.G.-M., C.-P.Y., I.E.-M., M.K. and T.L.; Visualization, J.G.-M.; Supervision, M.K. and T.L.; Project Administration, M.K. and T.L.; Funding Acquisition, M.K. and T.L.

## Material and Methods

### Key Resources Table

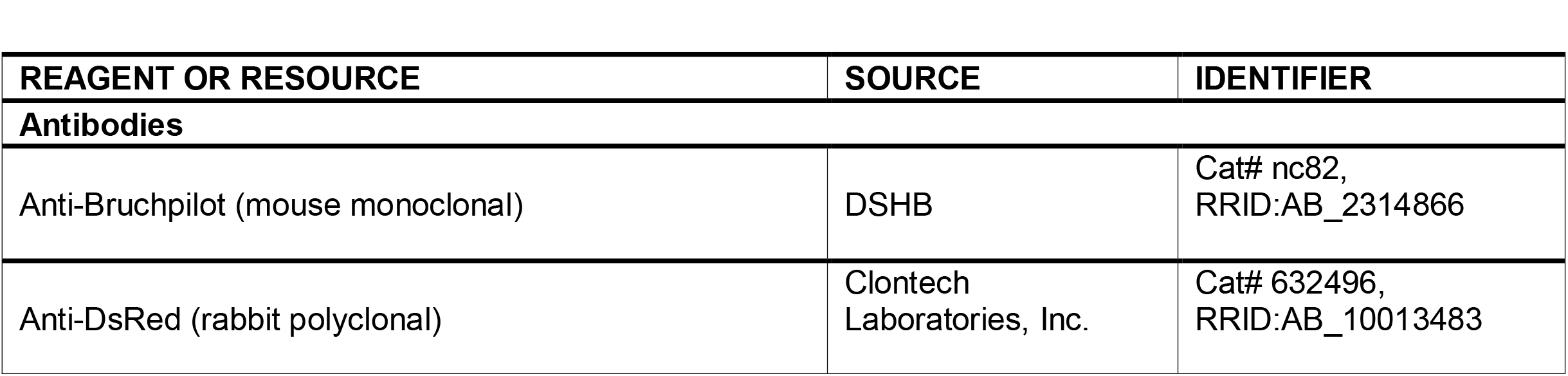
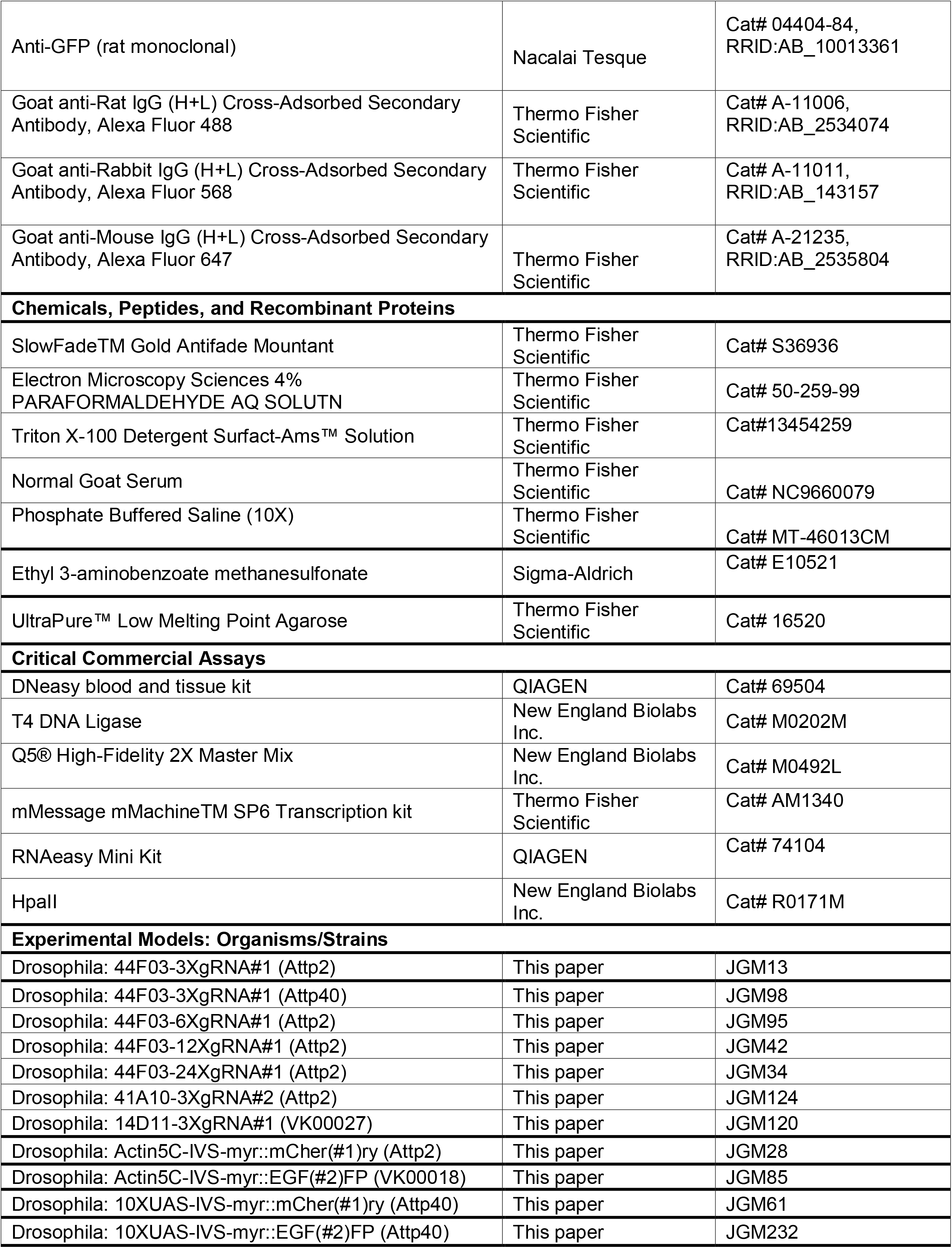
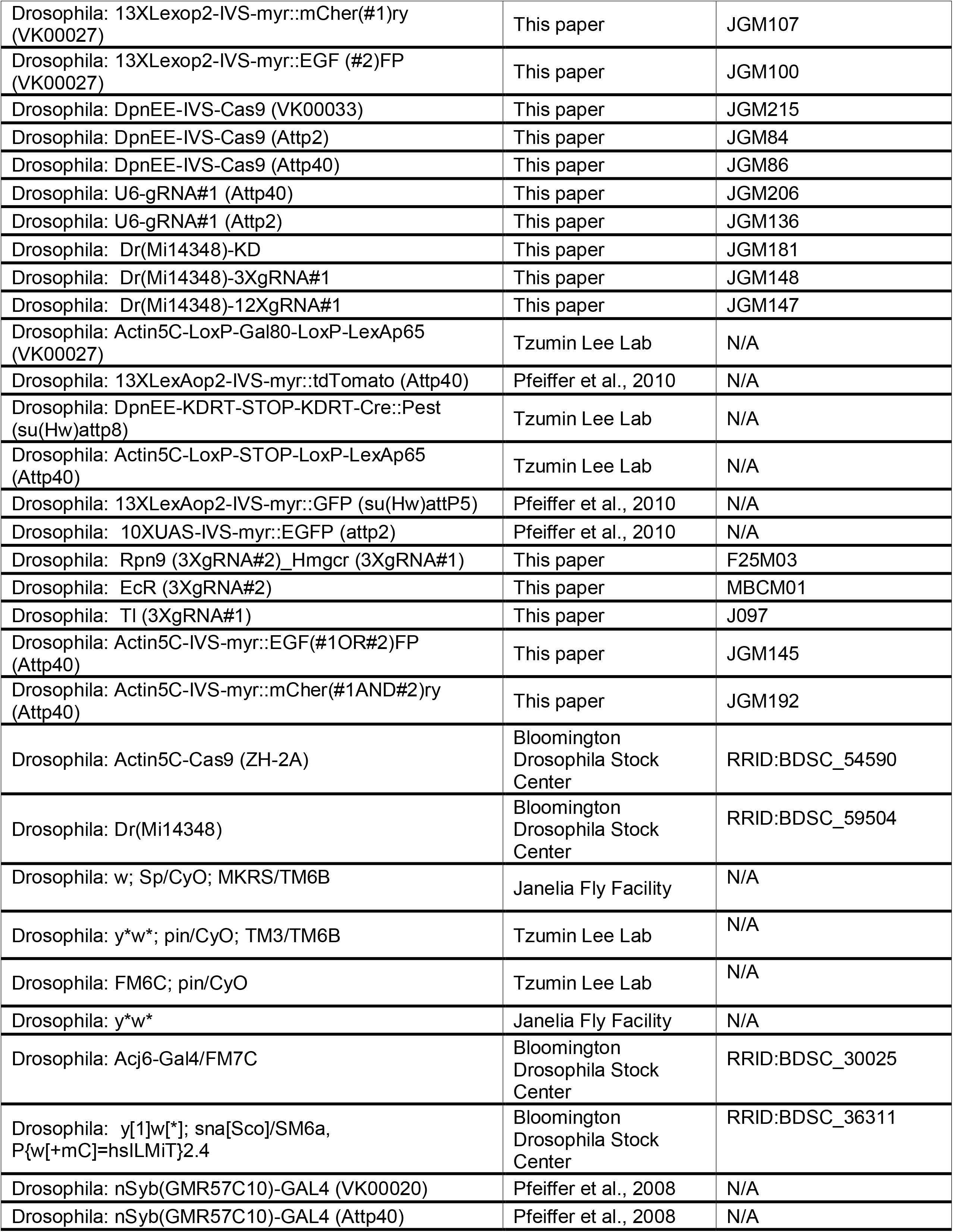
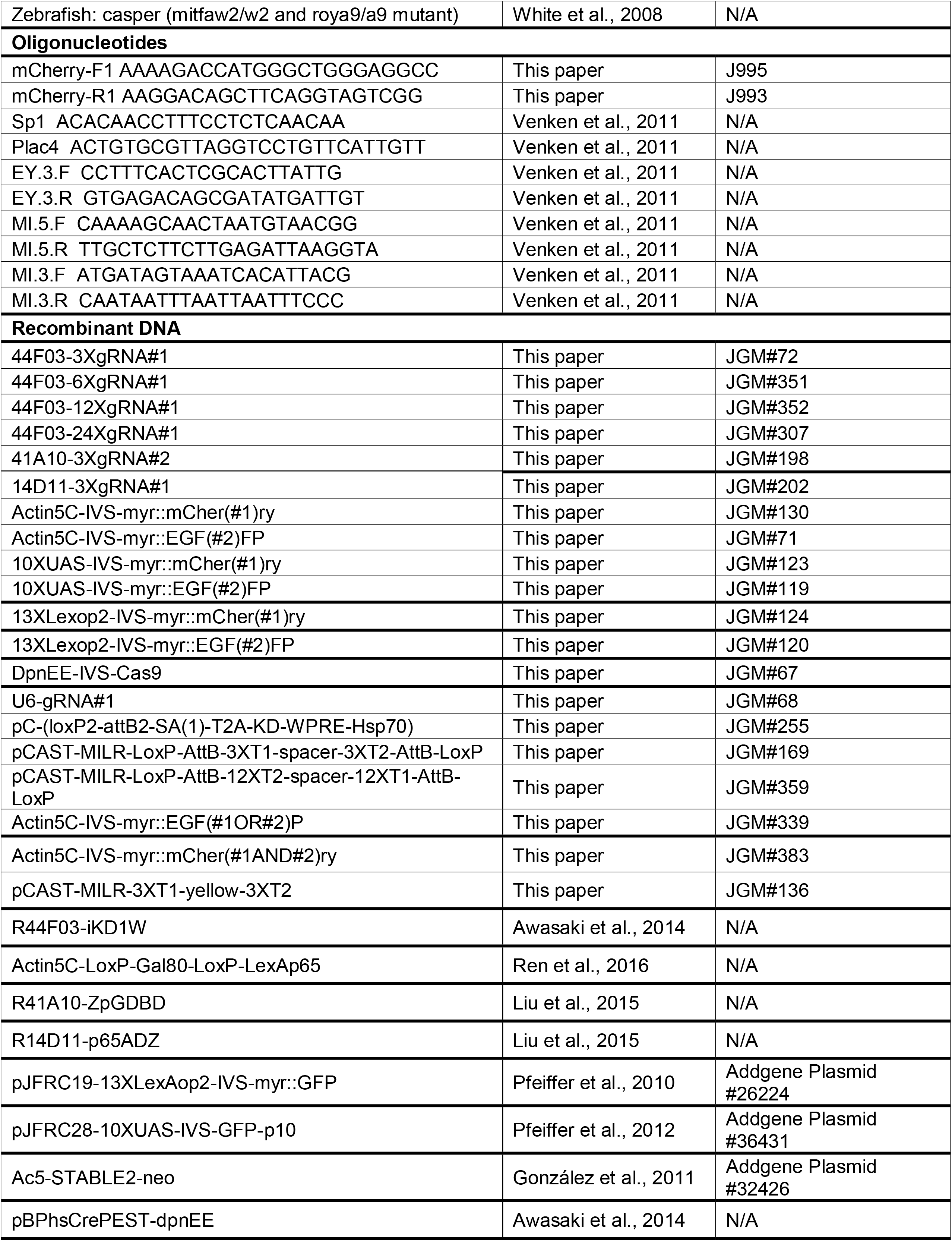
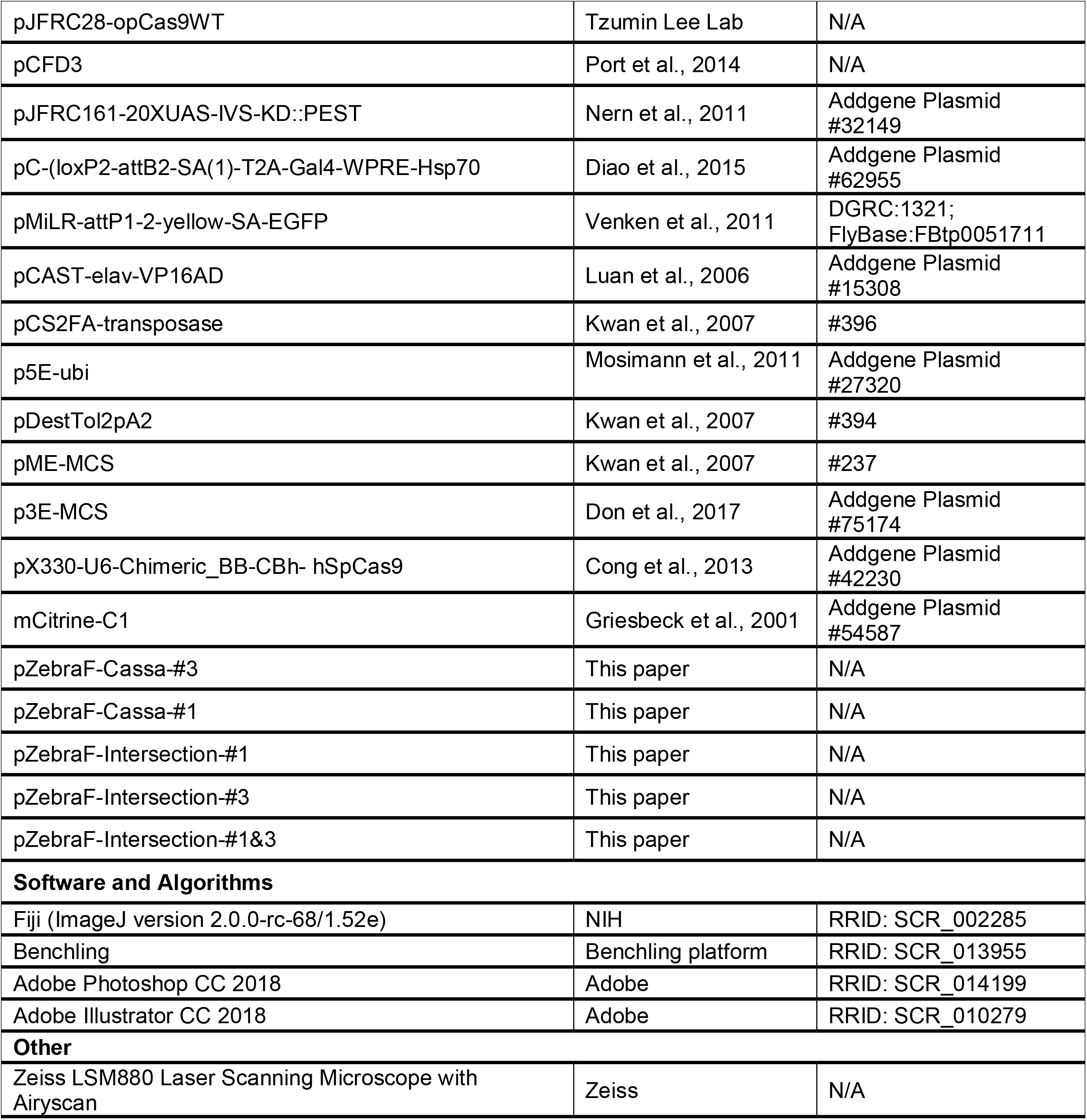

### Contact for Reagent and Resource Sharing

Further information and requests for resources and reagents should be directed to and will be fulfilled by the Lead Contact, Tzumin Lee.

### Experimental Models and Subject Details

#### Flies

Flies stocks were maintained in vials with standard fly food, in an incubator with constant temperature and humidity (50% relative humidity) under 12-hr light/dark cycles. Experimental crosses and the resulting progeny were maintained at 25°C under the same conditions.

#### Strains

The following stocks were used in this study: Actin5C-Cas9 (ZH-2A; BDSC#54590), nSyb(GMR57C10)-GAL4 (VK00020, Pfeiffer et al., 2008), Acj6-Gal4 (BDSC#30025), DpnEE-KDRT-STOP-KDRT-Cre::PEST (su(Hw)attp8; Tzumin Lee lab), Actin5C-LoxP-STOP-LoxP-LexAp65 (Attp40; Tzumin Lee lab), 13XLexAop2-myr::GFP (su(Hw)attp5; Pfeiffer et al., 2010), nSyb(GMR57C10)-GAL4(Pfeiffer et al., 2008), w; Sp/CyO; MKRS/TM6B (Janelia Fly Facility) and Dr(Mi14348) (BDSC#59504). In addition, we generated the following lines: 44F03-3XgRNA#1 (Attp2), 44F03-3XgRNA#1 (Attp40), 44F03-6XgRNA#1 (Attp2), 44F03-12XgRNA#1 (Attp2), 44F03-24XgRNA#1 (Attp2), 41A10-3XgRNA#2 (Attp2), 14D11-3XgRNA#1 (VK00027), Actin5C-IVS-myr::mCher(#1)ry (Attp2), Actin5C-IVS-myr::EGF(#2)FP (VK00018), 10XUAS-IVS-myr::mCher(#1)ry (Attp40), 10XUAS-IVS-myr::EGF(#2)FP (Attp40), 13XLexAop2-myr::mCher(#2)ry (VK00027), 13XLexAop2-myr::EGF(#2)FP (VK00027), DpnEE-IVS-Cas9 (VK00033), DpnEE-IVS-Cas9 (Attp2), DpnEE-IVS-Cas9 (Attp40), U6-gRNA#1 (Attp40), U6-gRNA#1 (Attp2), Dr(Mi14348)-KD, Dr(Mi14348)-3XgRNA#1, Dr(Mi14348)-12XgRNA#1. Actin5C-IVS-myr::EGF(#1OR#2)FP(Attp40) and Actin5C-IVS-myr::mCher(#1AND#2)ry (Attp40).

A list of the full genotypes for the flies used in each figure is in Table S1.

#### Generation of Transgenic Fly Lines

a. **Plasmid generation** Plasmids to generate transgenic fly lines were designed with Benchling (Benchling platform). To build these, we used standard molecular cloning techniques, including restriction digest/ligation and PCR assembly. The sequences for the different gRNAs were chosen by using Benchling (Benchling platform) based on their low off-target (Hsu et al., 2013) and their high on-target activity (Doench et al., 2014). The lowest score was 71.3 and 93.9 for on-target and off-target activity respectively. All inserts were verified by sequencing before injection:

1. 44F03-3XgRNA#1, 44F03-6XgRNA#1, 44F03-12XgRNA#1, 44F03-24XgRNA#1: we first generated 44F03-3XgRNA#1: KD in R44F03-iKD1W (Awasaki et al., 2014) was replaced (HindIII/KpnI) with a 3XgRNA#1 cassette (synthetized by Genscript). This 3XgRNA#1 cassette contains three copies of a gRNA#1 flanked by the ribozymes HammerHead and HDV (He et al., 2017). We then digested this plasmid to generate two fragments: KpnI-3XgRNA#1-SgrAI and AgeI-3XgRNA#1-HindIII. As SgrAI and AgeI produce compatible ends, both fragments were inserted into a KpnI/HindIII in the original plasmid. This reconstituted the original plasmid, now with a 6XgRNA#1 insert. This plasmid was used to repeat the process to generate 44F03-12XgRNA#1 and 44F03-24XgRNA#1.
2. 41A10-3XgRNA#2: a KpnI-3XgRNA#1-HindIII fragment (from 44F03-3XgRNA#1) was inserted into a KpnI/HindIII site in R41A10-ZpGDBD (Liu et al., 2015), thus replacing ZpGDBD.
3. 14D11-3XgRNA#1: a KpnI-3XgRNA#1-Hsp70Bb-SpeI fragment (from 44F03-3XgRNA#1) was inserted into a KpnI/SpeI site in R14D11-p65ADZ (Liu et al., 2015), thus replacing p65ADZ.
4. Actin5C-IVS-myr::EGF(#2)FP: first, we generated the following fragments by PCR from pJFRC19-13XLexAop2-IVS-myr::EGFP (Pfeiffer et al., 2010; Addgene #26224): a) KpnI-IVS-myr::EGFP(1-529bp)-2XgRNA#2-NheI, b) NheI-2XgRNA#2(inverted)-EGFP(130-717bp)-Hsp70Bb_Terminator-HindIII. We then cloned both fragments into a KpnI/HindIII site in Actin5C-LoxP-Gal80-LoxP-LexAp65 (Ren et al., 2016), thus replacing the loxP(opGal80)-Gal4 cassette.
5. 10XUAS-IVS-myr::EGF(#2)FP: a XhoI-myr::EGF(#2)FP-Hsp70Bb_Terminator-XbaI (from Actin5C-IVS-myr::EGF(#2)FP) fragment was inserted into pJFRC28-10XUAS-IVS-EGFP-p10 (Pfeiffer et al., 2012; Addgene#36431), thus replacing the EGFP.
6. 10XUAS-IVS-myr::mCher(#1)ry: we used Ac5-STABLE2-neo (González et al., 2011; Addgene #32426) for PCR amplification to generate the following fragments: a) BamHI-mCherry(1-528bp)-2XgRNA#1(inverted)-NheI, b) -NheI-2XgRNA#1-mCherry(129-529bp)-XbaI. Then, we clone it into 10XUAS-IVS-myr::EGF(#2)FP, thus removing EGF(#2)FP.
7. Actin5C-IVS-myr::mCher(#1)ry: we replaced myr::EGF(#2)FP in Actin5C-IVS-myr::EGF(#2)FP by inserting a fragment EagI-myr::mCher(#1)ry-EagI from 10XUAS-IVS-myr::mCher(#1)ry.
8. Actin5C-IVS-myr::EGF(#1OR#2)P: similar to Actin5C-IVS-myr::EGF(#2)FP, but in this case the fragment b) was amplified with sites 2XgRNA#1 instead.
9. Actin5C-IVS-myr::mCher(#1AND#2)ry: a fragment NotI-myr (1-202bp)-1XgRNA#2-MluI was generated by PCR amplification from Actin5C-IVS-myr::mCher(#1)ry. We also designed a synthetic gBlock(IDT) for a fragment MluI-1XgRNA#2-myr(3-253bp)-mCherry(1-528bp)-2XgRNA#1(inverted)-2XgRNA#1-mCherry(129-583)-XbaI. The final construct was then made by replacing myr::mCher(#1)ry in Actin5C-IVS-myr::mCher(#1)ry.
10. DpnEE-IVS-Cas9: a DpnEE driver was PCR amplified from pBPhsCrePEST-DpnEE (Awasaki et al., 2014) with sites HindIII and XhoI. We also generated a SpCas9 without extra nuclear localization signals by PCR amplification from pJFRC28-opCas9WT (a gift from Hui-Min Chen, Tzumin Lee lab). Both fragments were then cloned into a HindIII/XbaI site in pJFRC28-opCas9WT.
11. U6-gRNA#1: Annealed primers were cloned into a SapI site in pCFD3 (Port et al., 2014).
12. pC-(loxP2-attB2-SA(1)-T2A-KD-WPRE-Hsp70) (for RMCE): a T2A-KD fragment was amplified by PCR from pJFRC161-20XUAS-IVS-KD::PEST (Nern et al., 2011; Addgene#32140) and cloned in frame into a BamHI/PmeI site in pC-(loxP2-attB2-SA(1)-T2A-Gal4-WPRE-Hsp70) (Diao et al., 2015; Addgene#62955), thus replacing Gal4.
13. pCAST-MILR-3XT1-yellow-3XT2 (gene trap construct): a XmaI-3XgRNA#1-Hsp70Bb_terminator-SpeI cassette was PCR-amplified from 44F03-3XgRNA#1 and inserted into a XmaI/SpeI site in pMiLR-attP1-2-yellow-SA-EGFP (Venken et al., 2011; DGRC#1321), thus removing the SA-EGFP. Then, a 3XgRNA#2-Hsp70Bb was generated by PCR from 41A10-3XgRNA#2 and inserted (in reverse orientation) into a XbaI/XhoI site. Finally, a cassette containing the P-element inverted repeats was PCR-amplified from pCAST-elav-VP16AD (Luan et al., 2006; Addgene#15308) and cloned into a NheI site.
14. pCAST-MILR-LoxP-AttB-3XT1-spacer-3XT2-AttB-LoxP: First, we removed the marker yellow+ in pCAST-MILR-3XT1-yellow-3XT2 by inserting a small adapter in a SpeI/XhoI site. We then introduced a first AttB-LoxP cassette (gBlock, IDT) in XbaI/PacI. A second AttB-LoxP cassette (gBlock, IDT) was then introduced in BamHI/NotI.
15. pCAST-MILR-LoxP-AttB-12XT2-spacer-12XT1-AttB-LoxP: first, we generated a 12XgRNA#2 cassette following the same strategy explained above. Then, we cloned this and a 12XgRNA#1 cassette (from 44F03-12XgRNA#1, in reverse orientation) into a BamHI/KpnI site in pCAST-MILR-LoxP-AttB-3XT1-spacer-3XT2-AttB-LoxP.
b. **Fly injections** Constructs #1-11 contained an AttB site and were integrated into the corresponding landing sites using the PhiC31 system (Groth et al., 2004, Venken et al. 2006, Bischof et al. 2007). Constructs #12-14 contained two AttB sites and were integrated by RMCE (Bateman et al., 2006) into the MiMIC line Dr(Mi14348) (Venken et al., 2011). Construct #15 was randomly inserted into the genome by co-injection with the P-element transposase, generating 55 lines. Those lines were screened by crossing to y*w* and then balanced with FM6C; pin/CyO (for X chromosome insertions) or y*w*; pin/cyo; Tm3/TM6b (for 2^nd^ and 3^rd^ chromosome). All the embryo injections were performed by Rainbow Transgenic Flies Inc.
c. **Minos mobilization** Seven P-element insertions on the X chromosome (MAFF02, MNF03, MBQF02, MBQF022, MBQF01, F05F02, MAFF04) were remobilized to autosomes by crossing (G0 generation) to y[1]w[*]; sna[Sco]/SM6a, P{w[+mC]=hsILMiT}2.4, which expresses Minos transposase under heat-shock promoter regulation. Heat-shocks were performed in G0 larvae for 2h in a 37°C oven on 5 consecutive days. About 1500 G0 males with mosaic yellow+ body pattern were then collected and crossed (G1 generation) with y*w* virgin females, generating 115 yellow+ lines. These were balanced with y*w*; pin/CyO; TM3/TM6B.
d. **Mapping of insertion sites**Transposon insertions were mapped by inverse PCR and Sanger sequencing, following a modified protocol based on the one previously described (Venken et al., 2011; http://flypush.imgen.bcm.tmc.edu/pscreen/files/GDP_iPCRProtocol_051611.pdf). Genomic DNA was purified with a DNeasy Blood & Tissue Kit (Qiagen) and then digested with HpaII (NEB). After purification with a QIAquick Gel Extraction kit, digested DNA was ligated with T4 DNA Ligase (NEB) and used for two PCR reactions (one for each flanking region) with the Q5 polymerase (NEB) and the following primers: P-element: Sp1/Plac4 and EY.3.F/EY.3.R; b) Minos: MI.5.F/MI.5.R and MI.3.F/MI.3.R. We used the same primers to sequence the PCR product after gel extraction.

#### Zebrafish

All experiments were conducted according to the NIH guidelines for animal research and were approved by the Institutional Animal Care and Use Committee of Janelia Research Campus.

Animals were maintained at 28.5°C following standard procedures (Westerfield et al., 2000) and staged according to days post fertilization (dpf).

#### Strains

All the experiments were performed on *casper* background (White et al., 2008).

#### Injections

Adults (3 months to 2 years old,) were mated to produce embryos. Tol2 mRNA was synthetized from linearized plasmid using the mMessage mMachine SP6 Transcription kit (Thermo Fisher Scientific) and purified before injection into the zebrafish embryos using RNAeasy Mini Kit (Qiagen). About 400 embryos for each experiment were injected at 1-cell-state with 1-2 nanoliters of 25 ng/ul of Tol2 transposase mRNA and 25ng/ul of the corresponding Tol2 CaSSA plasmid. Fluorescence was examined from 1 dpf.

#### Plasmid generation

Injection plasmids were generated by multisite-Gateway cloning (Invitrogen) recombining the ubiquitous promoter containing vector p5E *ubi* (Mosimann et al., 2011; Addgene #27320), a pME vector carrying a triple cistron of SpCas9::P2A::mCherry::T2A::mCitrineCaSSAreporter followed by polyA signal, a p3E vector containing the corresponding U6:gRNA sequence and a pDestTol2 vector (Kwan et al., 2007).

The pME vectors were built as follows: first, we generated a fragment containing KpnI-SpCas9::T2A::mCitrineCaSSAreporter-polyA-SacII by Golden Gate cloning. KpnI-SpCas9 was amplified from pX330-U6-Chimeric_BB-CBh-hSpCas9 (Cong et al., 2013). For the mCitrineCaSSAreporter, mCitrine was amplified by PCR from mCitrine-C1 (Griesbeck et al., 2001), either with one (mCitr(#1)trine) or two switch cassettes (mCitr(#1)trin(#3)ine) interrupting the open reading frame. The first switch cassette contained a target site for the gRNA #1 located between two direct repeats (150-251bp of mCitrine). For the second switch cassette the target site was recognized by the gRNA #3 and the two direct repeats spanned 400-501 bp of the mCitrine sequence. Finally, pME-MCS (Tol2Kit #237) was digested by KpnI/SacII and ligated to KpnI-SpCas9::T2A::mCitrineCaSSAreporter-polyA-SacII. Subsequently, Gibson assembly (NEB) was used to clone a P2A-Cherry gBlock (IDT) between SpCas9 and the T2A::mCitrineCaSSAreporter (mCitr(#1)trine or mCitr(#1)trin(#3)ine), previously digested with FseI/MfeI.

The p3E vectors were built by digestion of the p3E-MCS (Addgene #75174) with SalI/BamHI to clone a U6c:gRNA#1 sequence or by SpeI/SacII to clone U6d:gRNA#3, or a combination of both (U6 promoters described in Yin et al., 2015). All U6:gRNA sequences were synthetized as gBlocks and cloned by Gibson assembly.

### Immunohistochemistry and confocal imaging

Brain tissues were dissected from larvae or adults in PBS and fixed with 4% paraformaldehyde (Thermo Fisher Scientific) in PBS for 40 minutes. After rinsing three times in 0.5% Triton X-100 (Thermo Fisher Scientific), the brains were incubated at 4°C overnight with primary antibodies diluted in PBS and 2% normal goat serum (Thermo Fisher Scientific). The primary antibodies used in this study were mouse anti-Bruchpilot (1:50; DSHB), rabbit anti-DsRed (1:500; Clontech) and rat anti-GFP (1:500; Nacalai). Next day, the brains were rinsed three times in PBS and transferred to a dilution of secondary antibodies in PBS. Secondary antibodies (1:1000; Thermo Fisher Scientific) included Alexa 488-conjugated goat anti-Rat IgG, Alexa 568-conjugated goat anti-Rabbit and Alexa 647 goat anti-Mouse IgG. After 4 hours at room temperature, the brains were washed again in PBS and mounted using SlowFade^TM^ Gold Antifade Mountant (Thermo Fisher Scientific).

For Zebrafish analyses, animals were anesthetized by bath application of 0.02% w/v solution of Ethyl 3-aminobenzoate methanesulfonate (Sigma-Aldrich, St. Louis) in filtered fish system water for 1 min and mounted in a drop of 1.6% low melting point agarose (Invitrogen) over a glass-bottomed dish.

Images were collected by a Zeiss LSM 880 confocal microscope and processed using Fiji(NIH) and Adobe Photoshop CC 2018 (Adobe).

### PCR from transgenic flies

To sequence the outcome of DNA repair events in those flies with a failed reconstitution of the Actin5C-mCher(#1)ry reporter (Figure S1), we first purified genomic DNA with a DNeasy Blood & Tissue Kit (Qiagen). We then performed a PCR reaction with the Q5 polymerase (NEB) and the primers mCherry-F1/mCherry-R1. Positive and negative controls were amplified in parallel producing the expected product or no product respectively.

### Quantification and Statistical Analysis

#### Assessing SSA effectiveness in single repair events

Female flies with Actin5C-Cas9 in the X chromosome were crossed (G0) to males with U6-gRNA#1 (2^nd^ chromosome) and the reporter Actin5C-mCher(#1)ry (3^rd^ chromosome) as shown in Figure S1. Then, we set up 100 individual G0 crosses to double balancer as follows: G0 males (50) were crossed to 3 double balancer virgin females and G0 females (50) were crossed to 2 double balancer males. To ensure that we quantified independent repair events, we randomly selected a single adult G1 fly (with the reporter and only one of the third chromosome balancers) from each vial and checked the expression of mCherry. We found a single mCherry expression pattern with strong and ubiquitous fluorescence. Finally we calculated the percentage of mCherry positive flies out of the total number of flies.

#### Gene trap analysis

Lines with transposon insertions were crossed to a line with Actin5C-Cas9 and two reporters, Actin5C-EGF(#2)FP and Actin5C-mCher(#1)ry. We then gave a semiquantitative score from 0 to 5 for each color, according to the extension of labeling. Subsequently, we calculated the percentage of lines with EGFP, mCherry, both reporters or no expression at all and express this as a percentage out of the total number of lines.

#### Effect of the gRNA copy number in the targeting effectiveness with the 44F03 driver

Transgenic lines with 3, 6, 12 or 24 copies of gRNA#1 under the regulation of the 44F03 driver were crossed to a reporter line with DpnEE-Cas9 (VK00033) and Actin5C-mCher(#1)ry. Then, we localized the Alad1, ALl1, ALv1, ALv2, EB and MB lineages, which were subjectively classified as complete or partial clones based on their size. We expressed this data as the percentage of hemibrains with complete or partial clones out of the total number for a minimum of 50 hemibrains. Note that for simplicity, we did not count the number of clonal units per hemibrain, which is 1 for all the lineages except for MB with 4 clonal units per hemibrain.

#### Effect of the gRNA copy number in the targeting effectiveness with the Dr driver

First, we identified the diverse lineages labeled after crossing Dr-3X or −12XgRNA#1 with flies bearing Actin5C-Cas9, Actin5C-mCher(#1)rry and Actin5C-EGF(#2)F. Then we calculated the percentage of hemibrains showing mCherry expression for each lineage out of a minimum of 60 hemibrains.

#### Testing the effectiveness of the intersection reporter

To quantify this parameter we first estimated the probability of MB (hemibrains) co-labeling with 41A10-gRNA#2 and 14D11-gRNA#1 when crossed to a line with Actin5C-IVS-myr::EGF(#2)FP, Actin5C-IVS-myr::mCher(#1)ry and DpnEE-IVS-Cas9. With this data we normalized the total number of hemibrains to the expected number considering the probability of co-labeling with both drivers. Then, we identified single MB clonal units in the intersection experiment and calculated the percentage of labeled MB clonal units out of the total number (corrected for the probability of co-labelling) of MB clonal units.

#### Zebrafish analyses

2 dpf positive fluorescent zebrafish larvae were randomly selected for imaging. We labeled and counted fluorescent cells using the Cell Counter plugin in the Fiji software (ImageJ version 2.0.0-rc-68/1.52e). A minimum of 16 cells per animal and 5 animals were analyzed in each experiment. The percentage of mCitrine positive cells out of the total number of mCherry positive cells was calculated.

## Supplemental Information

**Figure S1.**
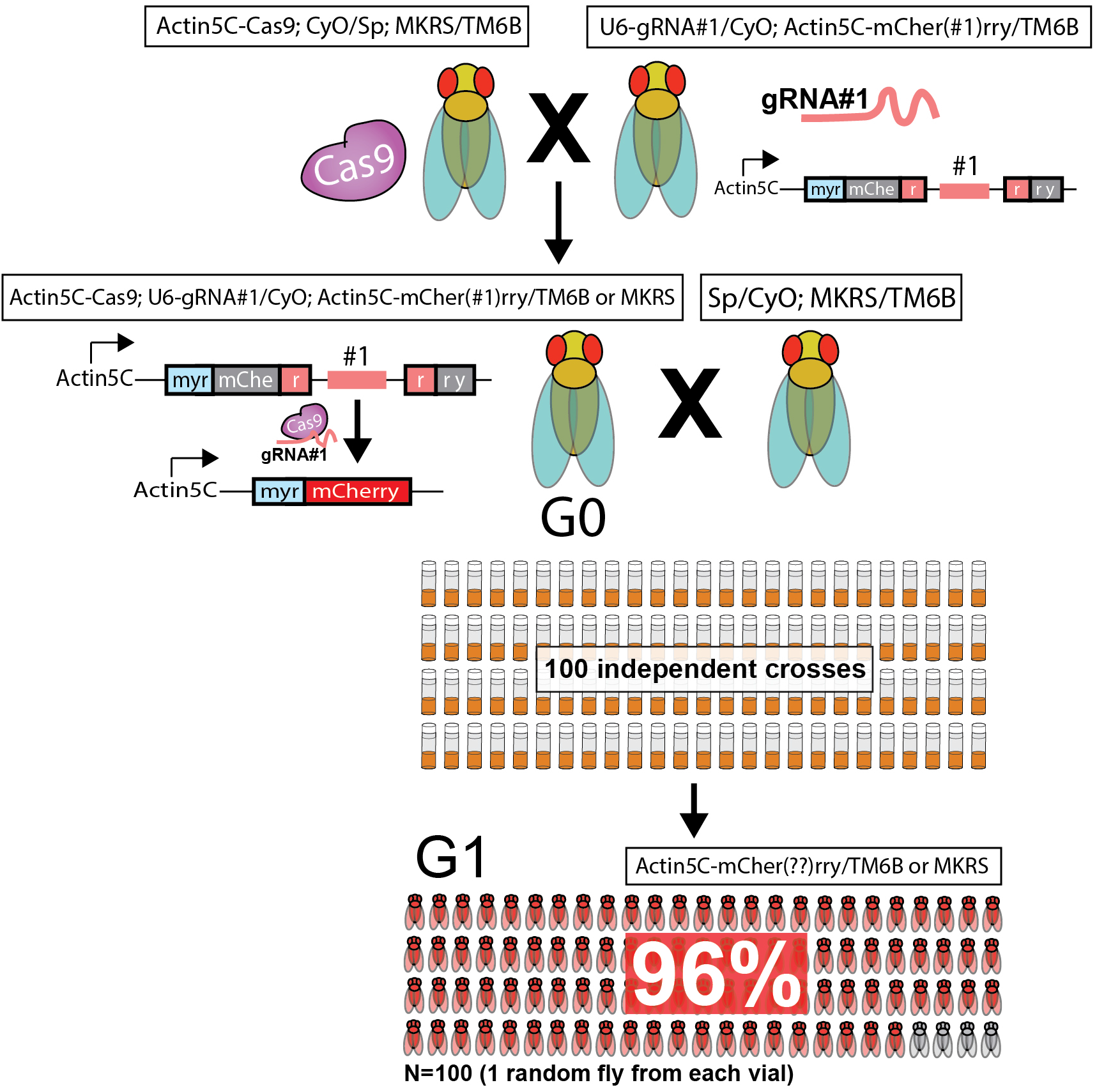
Effectiveness of SSA in independent DNA repair events, related to Figure 1. Flies bearing the three CaSSA components (G0) were generated by crossing virgin females with an Actin5C-Cas9 transgene to males containing both gRNA and reporter transgenes. To ensure that each repair event is independent, we set up 100 individual G0 crosses to double balancer in different vials. G0 males (50) were crossed to three double balancer virgin females and G0 females (50) were crossed to 2 double balancer males. Then, a single G1 adult fly bearing the reporter (with only one of the third chromosome balancers) was selected from each vial at random and checked for the expression of fluorescence (N=100 flies). Because Cas9 is driven by the ubiquitously expressed Actin5C promoter, the repair event could occur in any cell in the G0 fly. Only those events occurred in the germ line cells or their progenitors (i.e. pole cells) could be inherited (and therefore analyzed) by the G1 flies.

**Figure S2.**
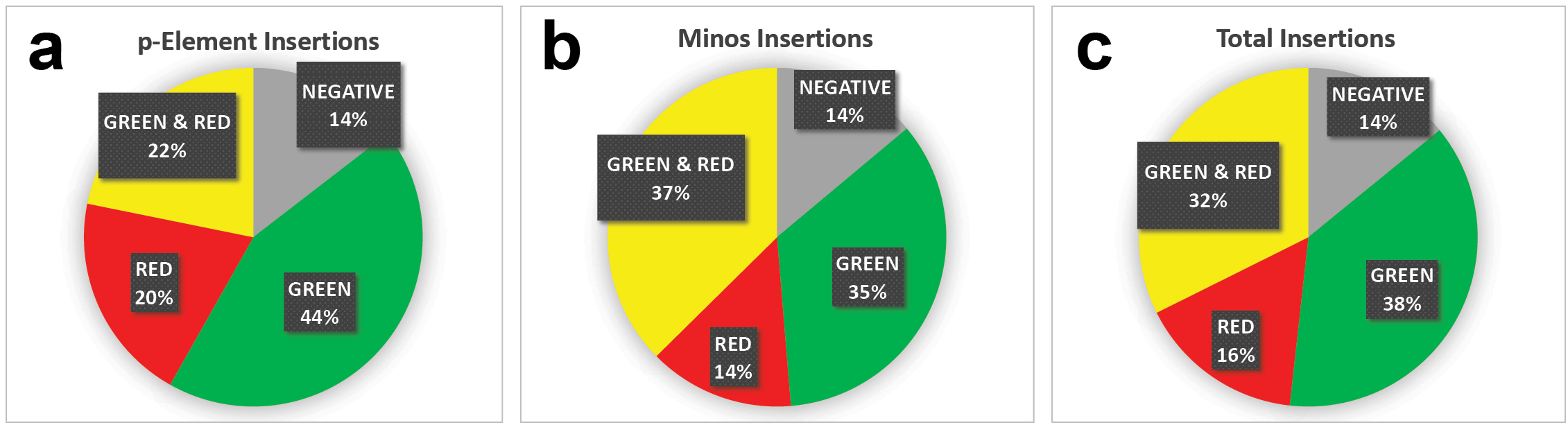
Gene trap efficiency, related to Figure 3. Gene trap constructs were inserted into the genome with either P-Element (55 insertions) or Minos (115 insertions) transposition. Percentage of P-element(A), Minos (B) or total gene trap insertions (C) exhibiting differential fluorescent reporter expression.

**Figure S3.**
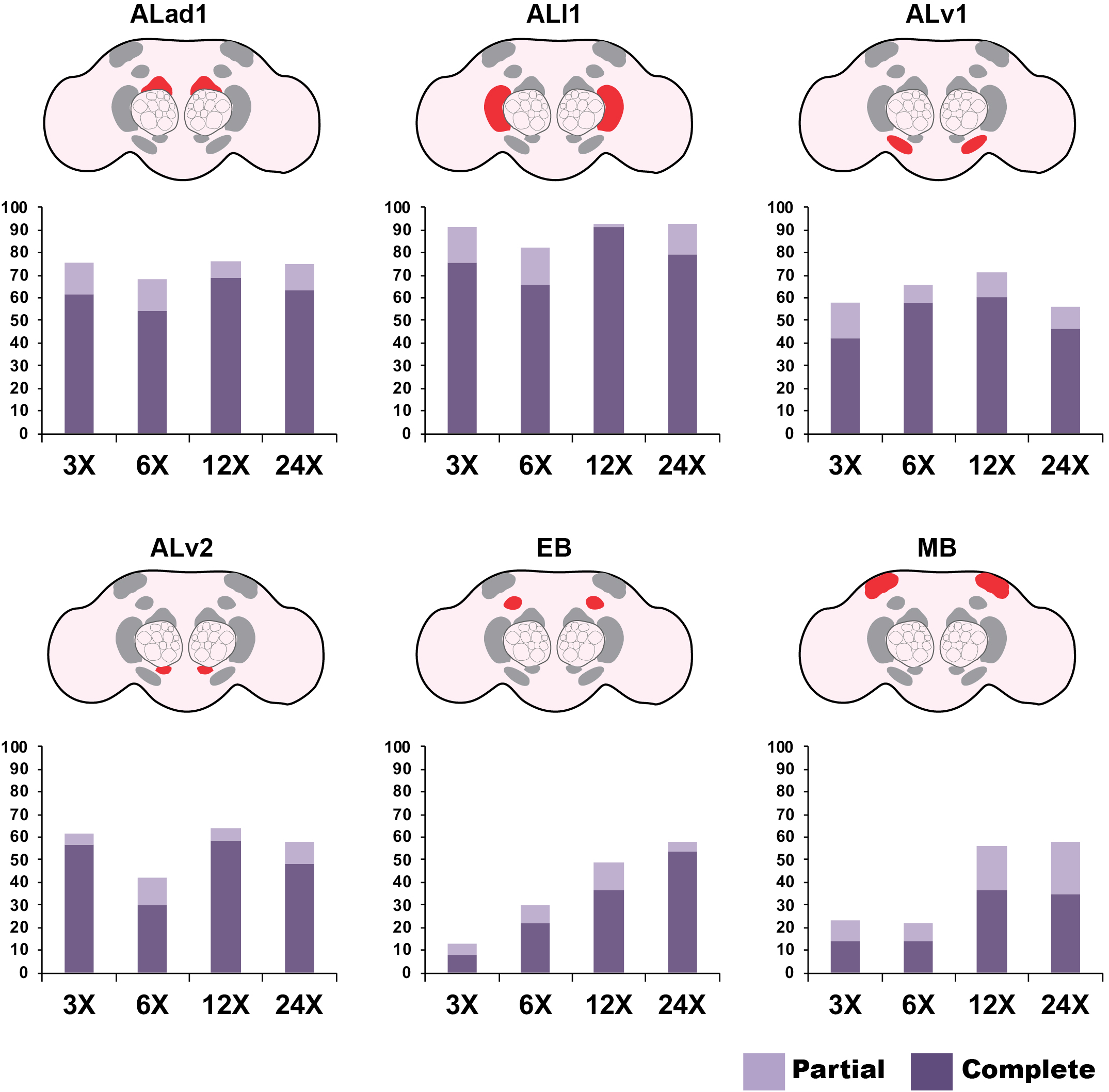
12 copies of gRNA are most effective to target lineages with the 44F03 driver, related to Figure 4. To enhance driver activity and minimize variability, increasing copy number of 44F03 driven gRNA was assessed. Six different lineages are targeted with the 44F03 driver, with varying levels of consistency. Each lineage was assessed individually for partial or complete coverage (see Material and Methods). The percentage of hemibrains with partial or complete coverage is represented for each lineage. For a majority of lineages, the highest percent of coverage is obtained with 12 copies of gRNA.

**Figure S4.**
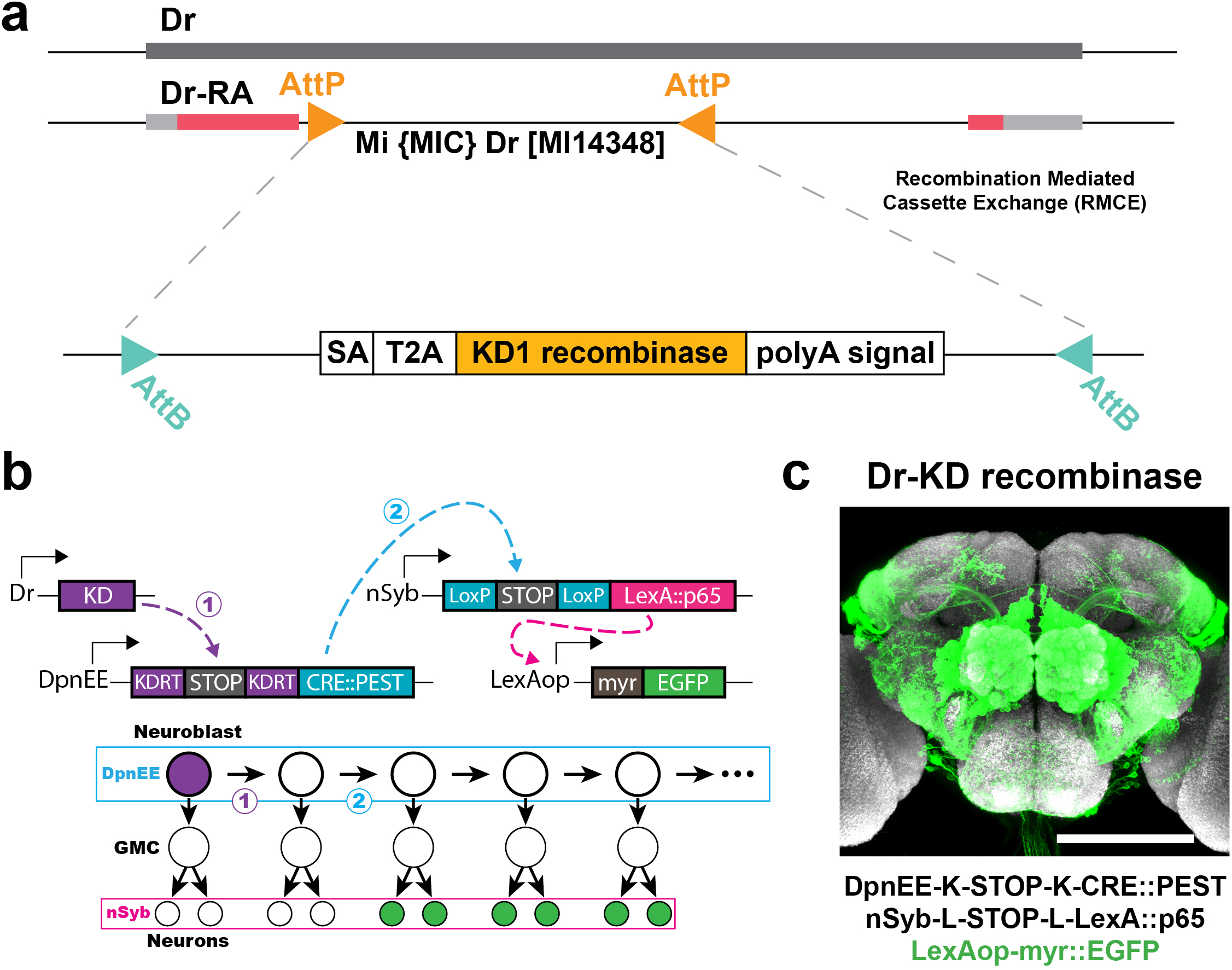
Dr pattern as revealed by driver immortalization with recombinases, related to Figure 4. A)Illustration showing the strategy to insert a KD recombinase in the Dr locus (Diao et al., 2015). In this case the cassette contains: 1) a splicing acceptor (SA) to modify the endogenous splicing so that our cassette is included in the mRNA, 2) a T2A peptide to express the recombinase from the same open reading frame as the endogenous gene, 3) the KD1 recombinase sequence and 4) a polyA signal, necessary for mRNA processing and stability. B)Schematics to illustrate the immortalization of the Dr driver in neuroblasts, based on a cascade of recombinases. The expression of KD (purple) results in the removal of a STOP cassette. This leads to the expression of Cre recombinase in those cells positive for DpnEE (neuroblasts). Cre is fused to PEST, a degradation signal that prevents Cre from being retained in GMCs or neurons. The expression of Cre removes another STOP cassette, leading to the expression of LexA:p65 in those cells expressing nSyb (post-mitotic neurons). LexAp65 then activates a myristoylated form of EGFP (green). C)Dr expression pattern as revealed by immortalization in neuroblasts. Maximal projection picture of a whole-mount brain is shown. Green, immunohistochemistry for EGFP. Gray, nc82 counterstain. Representative image of N=25 brains. K=KDRT. L=LoxP. Scale bar = 150 µm.

**Table S1.** List of fly genotypes used in this study.

|  | Genotype |
| --- | --- |
| 1B | +/+; Sp/CyO; Actin5C-IVS-myr::mCher(#1)ry (Attp2) / TM6B |
| 1B | +/+; Sp / CyO; U6-gRNA#1 (Attp2) / Actin5C-IVS-myr::mCher(#1)ry (Attp2) |
| 1B | +/+; Sp/CyO; Actin5C-IVS-myr::mCher(#1)ry (Attp2), DpnEE-IVS-Cas9 (VK00033) / TM6B |
| 1C | +/+; Sp/CyO; Actin5C-IVS-myr::mCher(#1)ry (Attp2), DpnEE-IVS-Cas9 (VK00033) / U6-gRNA#1 (Attp2) |
| S1 | Actin5C-Cas9 (ZH-2A); Sp/CyO; MKRS/TM6B |
| S1 | +/+; U6-gRNA#1 (Attp40) /CyO; Actin5C-IVS-myr::mCher(#1)ry (Attp2) / TM6B |
| S1 | +/+; Sp / CyO; MKRS / TM6B |
| 2B | +/+; Actin5C-IVS-myr::EGF(#2)FP (VK00018) / CyO; Actin5C-IVS-myr::mCher(#1)ry (Attp2), DpnEE-IVS-Cas9 (VK00033) / 44F03-3XgRNA#1 (Attp2) |
| 2B | +/+; Actin5C-IVS-myr::EGF(#2)FP (VK00018) / CyO; Actin5C-IVS-myr::mCher(#1)ry (Attp2), DpnEE-IVS-Cas9 (VK00033) / 41A10-3XgRNA#2 (Attp2) |
| 2B | +/+; Actin5C-IVS-myr::EGF(#2)FP (VK00018) / 44F03-3XgRNA#1 (Attp40); Actin5C-IVS-myr::mCher(#1)ry (Attp2), DpnEE-IVS-Cas9 (VK00033) / 41A10-3XgRNA#2 (Attp2) |
| 2C | +/+; nSyb(GMR57C10)-GAL4 (VK00020) / CyO; 10XUAS-IVS-myr::EGFP (attp2) / TM6B |
| 2C | Acj6-Gal4/FM7C; Sp/ CyO; 10XUAS-IVS-myr::EGFP (attp2) / + |
| 2D | +/+; 10XUAS-IVS-myr::mCher(#1)ry (Attp40) / nSyb(GMR57C10)-GAL4 (VK00020) ; 44F03-3XgRNA#1 (Attp2) / DpnEE-IVS-Cas9 (Attp2) |
| 2D | Acj6-Gal4/FM7C; 10XUAS-IVS-myr::mCher(#1)ry (Attp40) /Cy0 ; 44F03-3XgRNA#1 (Attp2) / DpnEE-IVS-Cas9 (Attp2) |
| 3B | Actin5C-Cas9 (ZH-2A); Actin5C-IVS-myr::EGF(#2)FP (VK00018) / CyO; Actin5C-IVS-myr::mCher(#1)ry (Attp2), DpnEE-IVS-Cas9 (VK00033) / Rpn9 (3XgRNA#2)_Hmgcr (3XgRNA#1) |
| 3B | Actin5C-Cas9 (ZH-2A); Actin5C-IVS-myr::EGF(#2)FP (VK00018) / CyO; Actin5C-IVS-myr::mCher(#1)ry (Attp2), DpnEE-IVS-Cas9 (VK00033) / EcR (3XgRNA#2) |
| 3B | Actin5C-Cas9 (ZH-2A); Actin5C-IVS-myr::EGF(#2)FP (VK00018) / CyO; Actin5C-IVS-myr::mCher(#1)ry (Attp2), DpnEE-IVS-Cas9 (VK00033) / TI (3XgRNA#1) |
| S2 | Insertions crossed to: Actin5C-Cas9 (ZH-2A); Actin5C-IVS-myr::EGF(#2)FP (VK00018) / CyO; Actin5C-IVS-myr::mCher(#1)ry (Attp2), DpnEE-IVS-Cas9 (VK00033) / TM6B |
| S3 | +/+; Actin5C-IVS-myr::EGF(#2)FP (VK00018) / CyO; Actin5C-IVS-myr::mCher(#1)ry (Attp2), DpnEE-IVS-Cas9 (VK00033) / 44F03-3XgRNA#1 (Attp2) |
| S3 | +/+; Actin5C-IVS-myr::EGF(#2)FP (VK00018) / CyO; Actin5C-IVS-myr::mCher(#1)ry (Attp2), DpnEE-IVS-Cas9 (VK00033) / 44F03-6XgRNA#1 (Attp2) |
| S3 | +/+; Actin5C-IVS-myr::EGF(#2)FP (VK00018) / CyO; Actin5C-IVS-myr::mCher(#1)ry (Attp2), DpnEE-IVS-Cas9 (VK00033) / 44F03-12XgRNA#1 (Attp2) |
| S3 | +/+; Actin5C-IVS-myr::EGF(#2)FP (VK00018) / CyO; Actin5C-IVS-myr::mCher(#1)ry (Attp2), DpnEE-IVS-Cas9 (VK00033) / 44F03-24XgRNA#1 (Attp2) |
| S4C | Dpn-KDRT-STOP-KDRT-Cre::Pest (su(Hw)attP8); Actin5C-LoxP-STOP-LoxP-LexA::p65 (Attp40), 13XLexAop-myr::GFP (su(Hw)attP5) / CyO; UAS-KD (attp2) / Dr(Mi14348)-KD |
| 4B | Actin5C-Cas9 (ZH-2A); Actin5C-IVS-myr::EGF(#2)FP (VK00018) / CyO; Actin5C-IVS-myr::mCher(#1)ry (Attp2), DpnEE-IVS-Cas9 (VK00033) / Dr(Mi14348)-3XgRNA#1 |
| 4B | Actin5C-Cas9 (ZH-2A); Actin5C-IVS-myr::EGF(#2)FP (VK00018) / CyO; Actin5C-IVS-myr::mCher(#1)ry (Attp2), DpnEE-IVS-Cas9 (VK00033) / Dr(Mi14348)-12XgRNA#1 |
| 4C | Actin5C-Cas9 (ZH-2A); Actin5C-IVS-myr::EGF(#2)FP (VK00018) / CyO; Actin5C-IVS-myr::mCher(#1)ry (Attp2), DpnEE-IVS-Cas9 (VK00033) / Dr(Mi14348)-3XgRNA#1 |
| 4C | Actin5C-Cas9 (ZH-2A); Actin5C-IVS-myr::EGF(#2)FP (VK00018) / CyO; Actin5C-IVS-myr::mCher(#1)ry (Attp2), DpnEE-IVS-Cas9 (VK00033) / Dr(Mi14348)-12XgRNA#1 |
| 5B | +/+; Actin5C-IVS-myr::EGF(#2)FP (VK00018) / CyO; Actin5C-IVS-myr::mCher(#1)ry (Attp2), DpnEE-IVS-Cas9 (VK00033) / Dr(Mi14348)-12XgRNA#1 |
| 5B | +/+; Actin5C-IVS-myr::EGF(#2)FP (VK00018) / CyO; Actin5C-IVS-myr::mCher(#1)ry (Attp2), DpnEE-IVS-Cas9 (VK00033) / 41A10-3XgRNA#2 (Attp2) |
| 5B | +/+; Actin5C-IVS-myr::EGF(#2)FP (VK00018) / CyO; Actin5C-IVS-myr::mCher(#1)ry (Attp2), DpnEE-IVS-Cas9 (VK00033) / 14D11-3XgRNA#1 (VK00027) |
| 5B | +/+; Actin5C-IVS-myr::EGF(#1OR#2)P (Attp40) / DpnEE-IVS-Cas9 (Attp40); Dr(Mi14348)-12XgRNA#1 / 41A10-3XgRNA#2 (Attp2) |
| 5B | +/+; Actin5C-IVS-myr::mCher(#1AND#2)ry (Attp40) / DpnEE-IVS-Cas9 (Attp40); 14D11-3XgRNA#1 (VK00027), 41A10-3XgRNA#2 (Attp2) / TM6B |

**Table S2.**
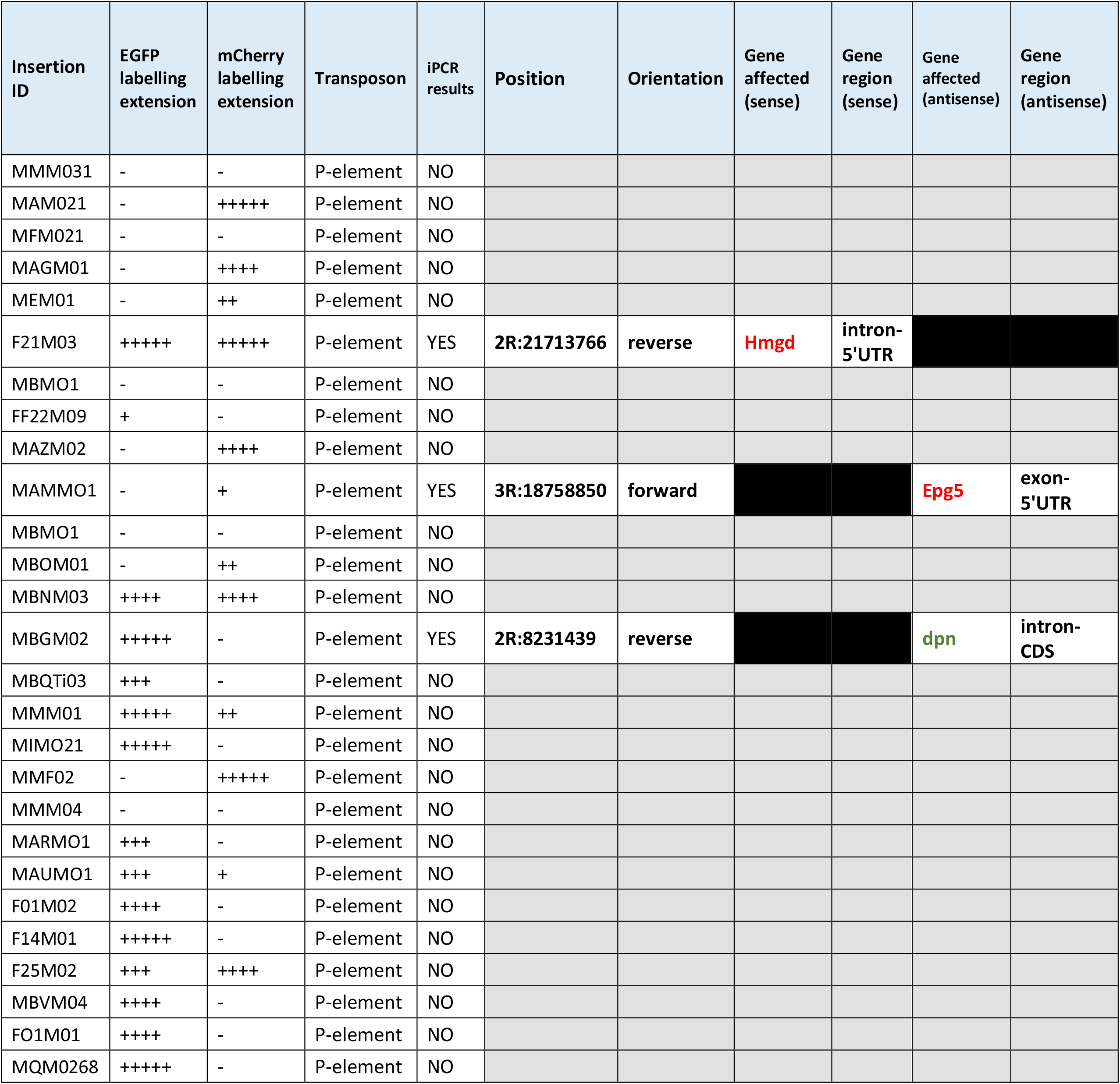
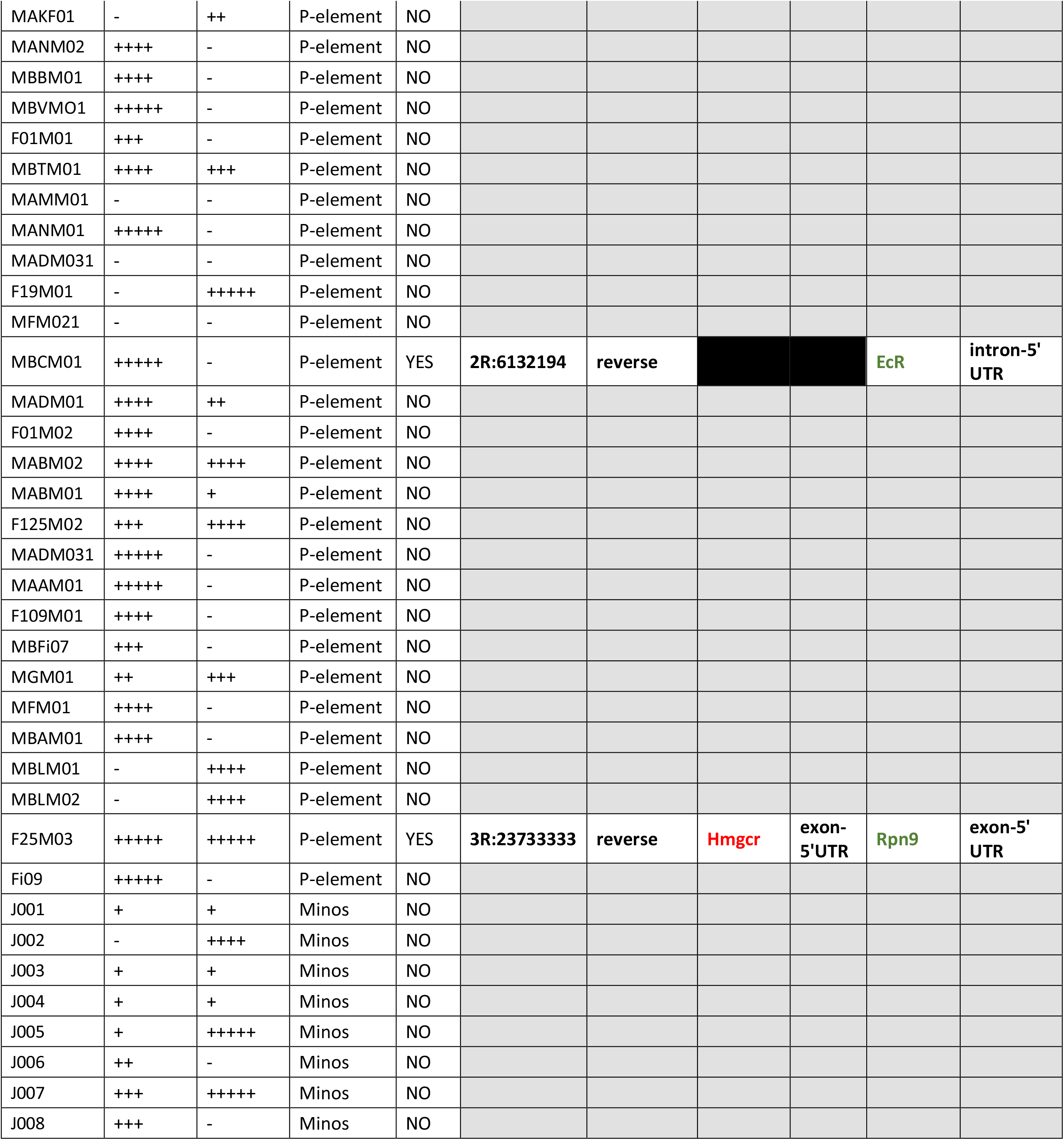
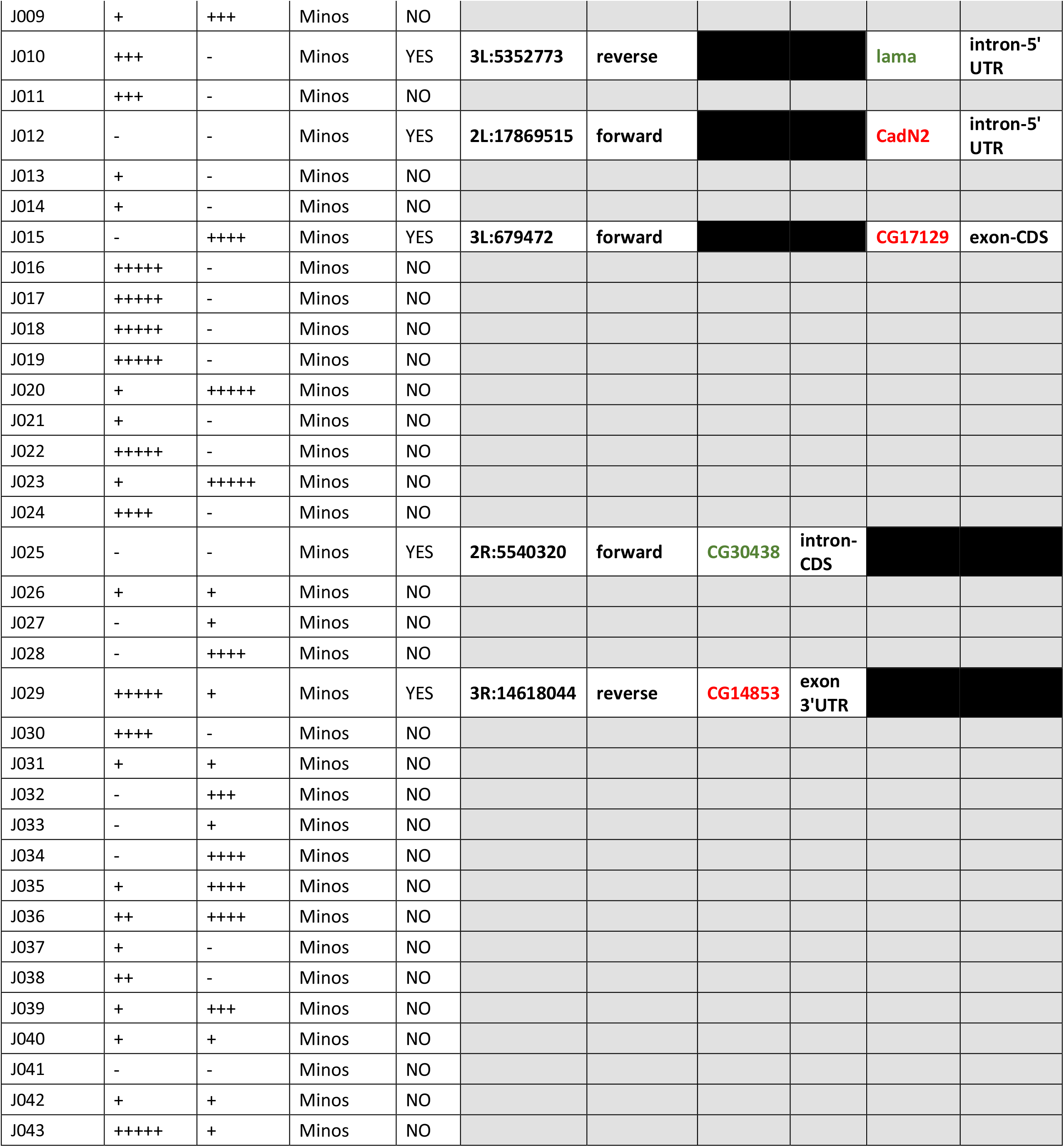
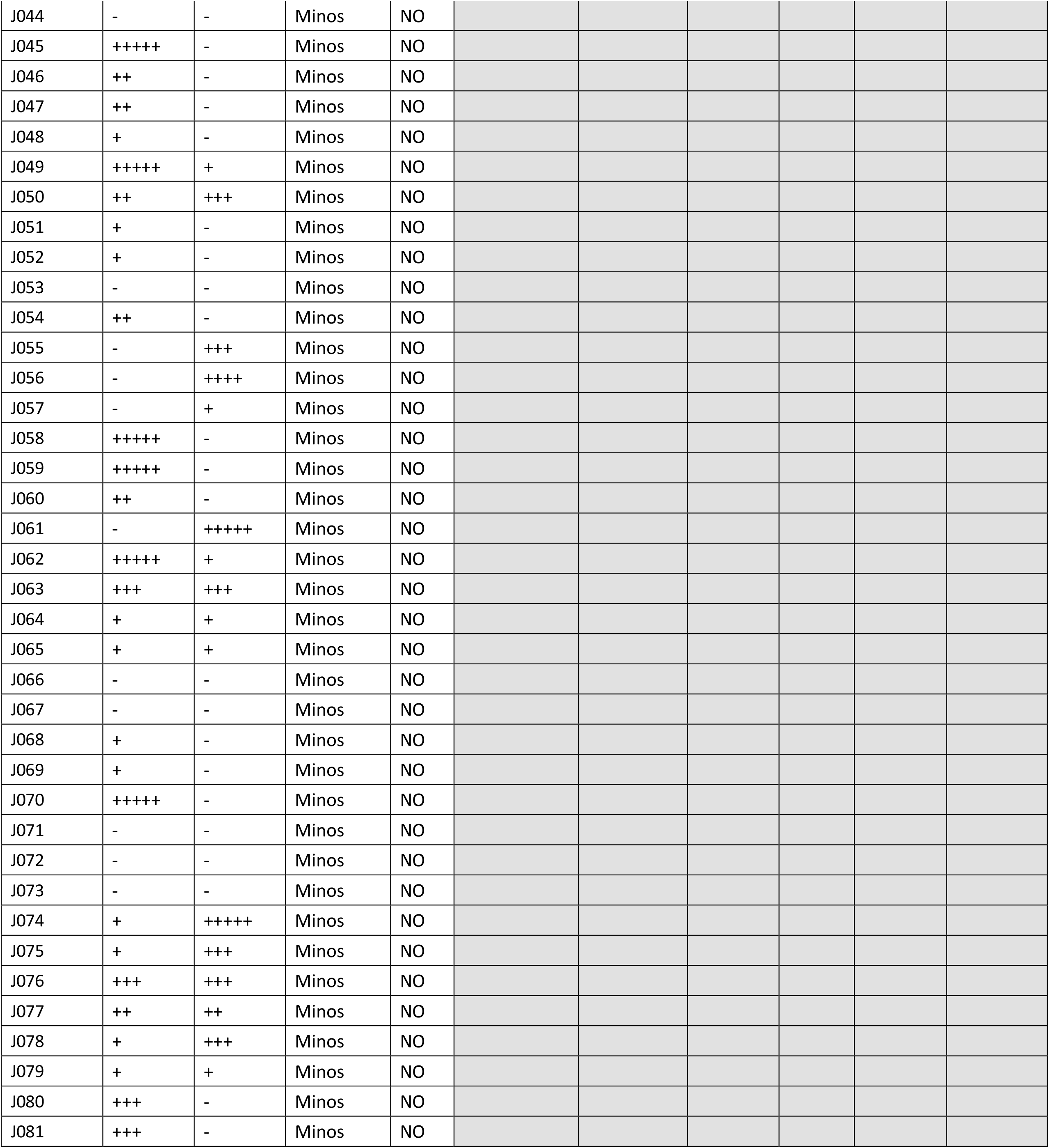
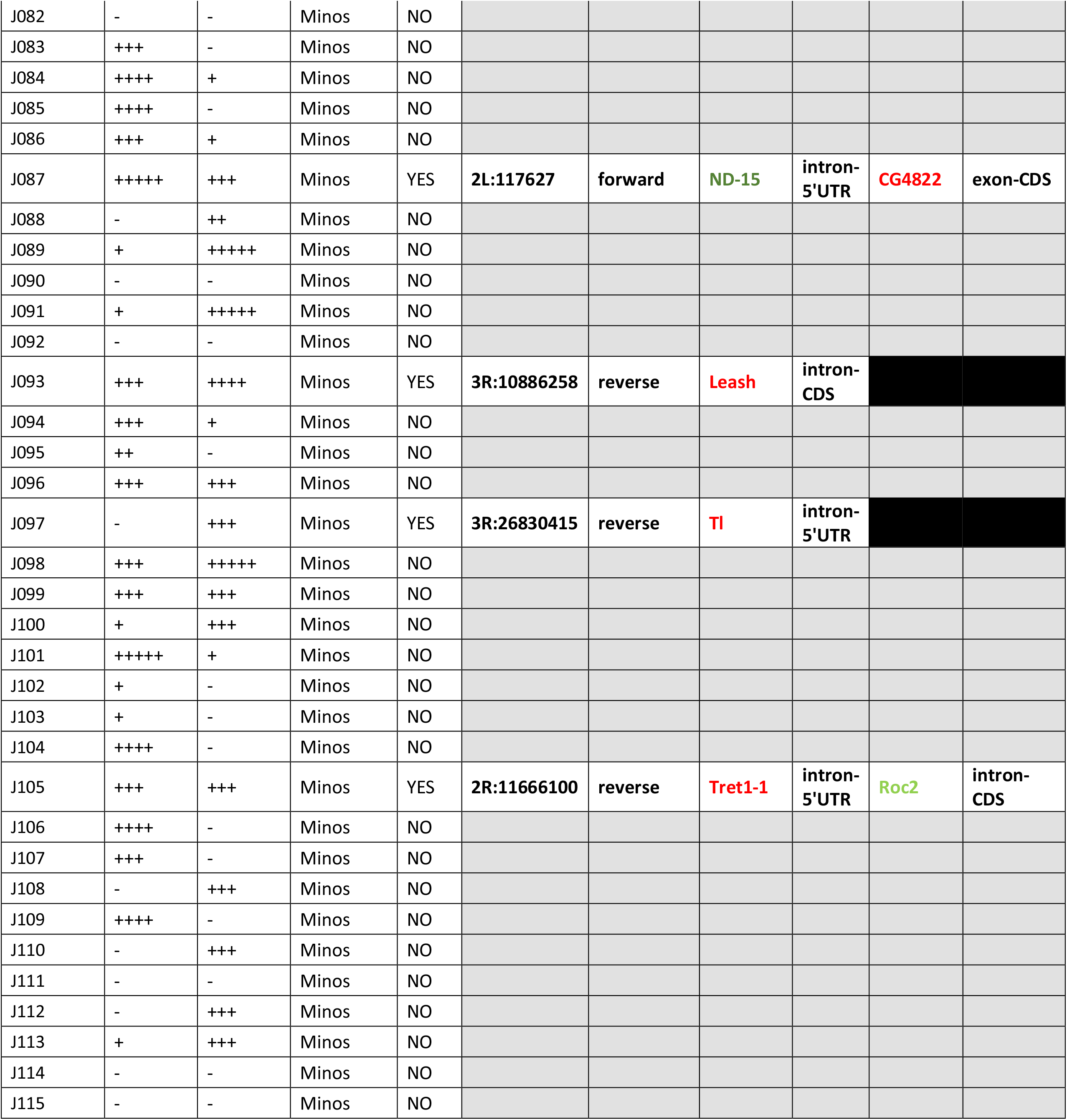
List of lines generated in the gene trap experiment.

